# EGFR-targeted antisense oligonucleotides modified with boron clusters offer an innovative approach to cancer chemo-radiotherapy

**DOI:** 10.1101/2025.08.07.668562

**Authors:** Damian Kaniowski, Katarzyna Ebenryter-Olbińska, Justyna Suwara, Katarzyna Kulik, Jeremy Hall, Dongfang Wang, Elaine Y. Kang, Rafał Dolot, Alessandro Somenzi, Agata Jakóbik-Kolon, Nicoletta Protti, Saverio Altieri, Marcin Kortylewski, Barbara Nawrot

**Affiliations:** Department of Immuno-Oncology, Beckman Research Institute at City of Hope Comprehensive Cancer Center, Duarte, CA 91010, USA; Centre of Molecular and Macromolecular Studies, Polish Academy of Sciences, Sienkiewicza 112, 90-363 Lodz, Poland; Department of Physics, Applied Nuclear Energy Laboratory (L.E.N.A.), University of Pavia, via Bassi 6, 27-100 Pavia, Italy; INFN—Istituto Nazionale di Fisica Nucleare, Sezione di Pavia, 27100 Pavia, Italy; Department of Inorganic, Analytical Chemistry and Electrochemistry, Faculty of Chemistry, Silesian University of Technology, Krzywoustego 6, 44-100 Gliwice, Poland

**Keywords:** Boron cluster, antisense oligonucleotide therapy, EGFR, drug delivery, immunotherapy, BNCT, XRT, crystallography

## Abstract

Suppression of cancer-associated EGFR expression by antisense oligonucleotides is an attractive strategy to augment the efficacy of radiotherapy in cancer treatment. Boron clusters act as selective ligands for the epidermal growth factor receptor (EGFR) and provide an innovative platform for the delivery of naked boron cluster-conjugated therapeutic nucleic acids to cancer cells. Here, we demonstrate that the novel inhibitor B-ASO^LNA^-CHOL can also enhance efficient and cancer cell-selective uptake via the low-density lipoprotein receptor (LDLR). This conjugate exhibit strong cytotoxic effects on skin and liver cancer cells in combination with BNCT or XRT. We confirmed that intratumoral injections of B-ASO^LNA^-CHOL combined with local XRT significantly reduced tumor size and EGFR expression in human A431 xenografts implanted in immunodeficient mice. Our findings suggest that B-ASO-CHOL containing a CpG ODN motif exhibits immune adjuvant properties. The results underscore the potential of B-ASO^LNA^-CHOL technology for therapeutic applications in radiation immuno-oncology.

Graphical abstract

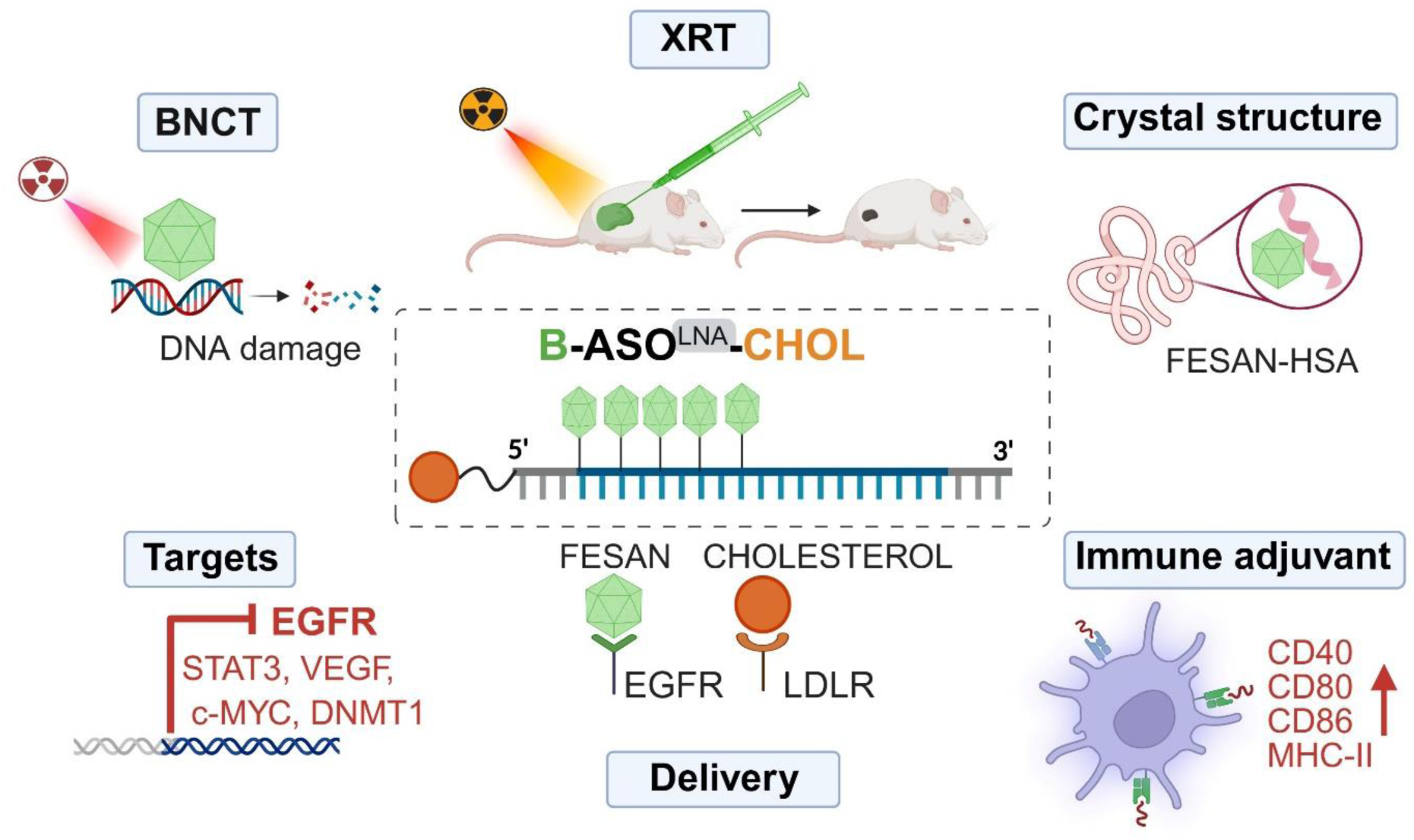

## Introduction

Boron neutron capture therapy (BNCT) is an internal binary radiotherapy strategy that relies on the selective accumulation of boron-10 (^10^B) isotope-labelled compounds in cancer cells. Nonradioactive ^10^B is exposed to epithermal neutrons to generate two high-energy particles (^4^He and ^7^Li) nucleus that kill cancer cells within a diameter of 4–10 μm, while neighboring normal cells survive.^1^ For BNCT to achieve a sufficient therapeutic effect, the tumor uptake of boron atoms should be in the range of 20-50 μg ^10^B/g cells. Despite extensive efforts in the field of BNCT, the development of more effective boron-containing compounds that can also serve as potent tumor growth inhibitors remains an ongoing need.^2^

The world’s first BNCT drug, Steboronine® (Borofalan), was approved by the Japanese Ministry of Health, Labor and Welfare on March 25, 2020 for the treatment of unresectable, locally recurrent or locally advanced head and neck cancer.^3^ The development of new boron-containing agents for BNCT must meet certain criteria, including a cancer-to-normal cell selectivity ratio (C/N) of at least 2.5:1, high accumulation of boron-rich compounds in the tumor without removal, and low cytotoxicity.^4^

Boron clusters are promising small molecules for their potential use in BNCT, especially FESAN ([Fe(C_2_B_9_H_11_)_2_]^-^), which can also serve as a radiosensitizer in cancer treatment. Radiation therapy (XRT) is a standard and widely employed external treatment modality for cancer; however, tumor resistance to XRT remains a major challenge, often reducing its therapeutic effectiveness.^5^ One strategy to overcome this limitation is the use of radiosensitizers, which increase tumor sensitivity while sparing healthy tissue. Combining radiotherapy with immunotherapy also offers a promising approach to improve cancer outcomes.^6^

A variety of different cancers express increased levels of the epidermal growth factor receptor (EGFR), which is a constant challenge in oncology. Hyperactivation of EGFR leads to enhanced cancer cells proliferation, migration, invasion, angiogenesis and chemoresistance.^7^ Moreover, EGFR can activate signal transducer and activator of transcription-3 (STAT3) which promotes tumor cell survival and mediates tumor-induced immunosuppression at multiple levels.^8,9^ Blockade of STAT3 synergizes with other promising immunotherapeutic agents, such as CpG oligodeoxynucleotides (CpG ODNs) and cancer vaccines, which promote Toll-like receptor 9 (TLR9) activation in immune cells.^10^ CpG ODNs are single-stranded synthetic compounds that contain one or more unmethylated cytosine-guanine (CG) dinucleotides within specific base-pair sequences known as CpG motifs. These ODNs exhibit strong immunoadjuvant properties and act as booster for anticancer agents in clinical trials.^11^

There are several FDA-approved EGFR inhibitors. Monoclonal antibodies (e.g., cetuximab, panitumumab) target the extracellular domain of EGFR, blocking ligand binding, while tyrosine kinase inhibitors (e.g., gefitinib, erlotinib, afatinib) inhibit its intracellular domain, thereby disrupting downstream signalling pathways essential for cancer cell proliferation. However, the durability of clinical response to these treatments is often limited or associated with significant side effects.^12^ This underscores the urgent need for better anti-EGFR molecular strategies to overcome the multistep resistance of EGFR^high^ human cancers.^13^

One potential approach involves the use of antisense oligonucleotides (ASOs), synthetic single-stranded DNA fragments, typically 13 to 25 nucleotides in length that can modulate oncogene expression. The typical ASO mechanism involves recruitment of RNase H to the DNA/RNA heteroduplex formed between the ASO and target mRNA, leading to cleavage of the RNA strand within the duplex.^14^ There are currently five FDA-approved ASOs based on the RNase H mechanism in clinical use: mipomersen (Kynamro, discontinued), inotersen (Tegsedi), volanesorsen (Waylivra), tofersen (Qalsody), and eplontersen (Wainua).^15^ We recently described an EGFR-targeted ASO highly modified with FESAN-boron clusters (B-ASO), enabling multifunctional therapeutic effects against cancer.^16^

Here, we developed a novel B-ASO^LNA^-CHOL (**11**) inhibitor that effectively reduces EGFR expression in cancers while enhancing BNCT- and XRT-dependent antitumor effects. We also identified a unique delivery mechanism for B-ASO^LNA^-CHOL agents (**11**), mediated by two receptors: EGFR and LDLR. As either a monotherapy or in combination with radiation, B-ASO^LNA^-CHOL demonstrated a direct killing effect in both in vitro and A431 xenograft mouse models. These findings support further preclinical investigation of B-ASO^LNA^-CHOL (**11**) in combination with radiation therapies.

## Results

### Chemical synthesis of an ASO EGFR conjugated with dual ligands: FESAN and a cholesterol moiety

To improve anticancer activity of the B-ASO conjugates, we modified the design to include the FESAN-boron cage (containing the natural abundance of ^10^B /^11^B isotopes) together with a cholesterol moiety (CHOL) to ensure effective cellular uptake using two synthetic steps. The first is the synthesis of the alkyne-functionalized DNA oligonucleotides, 25 nucleotides in length, using a solid-phase phosphoramidite method. Depending on the reagents used in “Step 1” (Figure 1A), different modifications were introduced into the growing oligonucleotide chain. For introduction of the CHOL moiety at the 3’-end of the oligonucleotide, the 3’-cholesteryl-TEG-CPG (CHOL-TEG CPG) was used instead of LCA CPG (long chain alkylamine controlled pore glass beads). To enable further modification of the oligonucleotides, appropriate phosphoramidite monomers were employed: cholesterol - CHOL(p), locked analog - LA(p), and 6-carboxyfluorescein - 6-FAM(p) monomers. This allowed the synthesis of compounds **2**, **4**, **6**, **12,** and **14** (abbr. U_PR_-ASO, U_PR_-ASO-CHOL, U_PR_-ASO-2CHOL, B-ASO^FAM^ and ASO^FAM^), respectively. To incorporate LNA modifications at the 5’-ends of the selected sequences (**8** and **10**, abbr. U_PR_-ASO^LNA^ and U_PR_-ASO^LNA^-CHOL), the oligonucleotide chain at the 5’-end was extended by three additional nucleotides (AAC) complementary to the target EGFR mRNA. In addition, a universal support (US CPG) was required to synthesize the LNA-3’-ends of these oligonucleotides. In the second step (”Step 2” in Figure 1A), boron clusters [alkyl azide derivative of (3,3′-iron-1,2,1′,2′-dicarbollide)^−1^, N_3_-alkyl-FESAN]^17^ were attached to the U_PR_ moieties post-synthetically. Conjugation was achieved using the Cu(I)-catalyzed [3+2] azide-alkyne Huisgen cycloaddition (CuAAC) reaction, a so-called click reaction. The final products **3**, **9**, **12** (B-ASO, B-ASO^LNA^ and B-ASO^FAM^, respectively), as shown in Figure 1A “Step 3”, were isolated from the reaction mixtures standard procedure (HPLC), while for compounds containing cholesterol units (B-ASO-CHOL **5**, B-ASO-2CHOL **7** and B-ASO^LNA^-CHOL **11**) a different approach based on Sep-Pak C18 cartridges was used (the HPLC profile for **11** is shown in Figure 1C). All compounds were analyzed by electrospray ionization quadrupole time-of-flight mass spectrometry (ESI-Q-TOF MS) (Figure S1). Cryo-TEM confirmed that B-ASO contains an oligonucleotide fragment with a FESAN rich-motif chain (Figure S2A).

**Figure 1.**
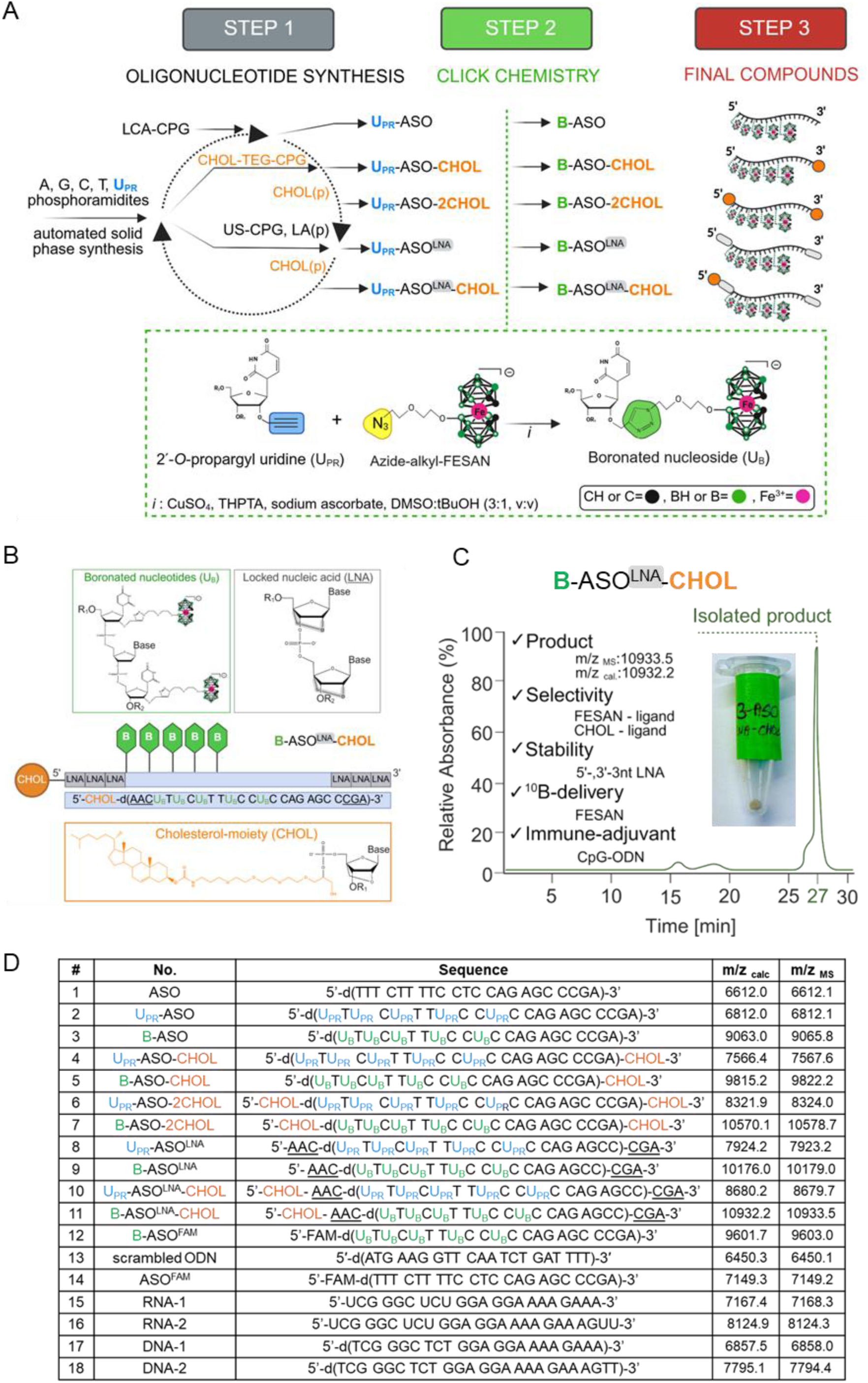
The design and synthesis of antisense oligonucleotides modified with a boron cluster and a cholesterol moiety (B-ASO^LNA^-CHOL). (**A**) UPR (blue) and B (green) stand for 2’-*O*-propargyluridine or boron cluster [(3,3’-iron-1,2,1’,2’-dicarbollide)^-1^, FESAN] (19% ^10^B and 81% ^11^B), CHOL (orange) is an abbreviation derived from the cholesterol moiety and LNA (grey) stands for locked analogs. Step 1 involves an automated solid phase oligonucleotide synthesis using commercially available supports (LCA CPG, CHOL-TEG-CPG and US CPG) and phosphoramidites (A, G, C, T, UPR, CHOL(p), LA(p)), followed by step 2 - CuAAC reactions between the corresponding propargylic derivative (blue label) of the oligonucleotide and the alkyl azide derivative of FESAN (yellow label) (the detailed reaction conditions are presented in part B) leading to the desired group of final compounds in step 3; (**B**) Schematic structure and sequence of B-ASO^LNA^-CHOL (**11**); (**C**) Profile of the preparative scale synthesis of B-ASO^LNA^-CHOL and its isolation by high performance liquid chromatography (HPLC). (**D**) Table of the tested oligonucleotides.

### Boron cluster (FESAN) brings unique properties to B-ASO^LNA^-CHOL

The downregulation of EGFR by B-ASO^LNA^ relies on RNase H (Figure 2A). To measure RNase H activity^14^, we compared the efficacy of B-ASO^LNA^ (**9**) in the duplex with a complementary 5’-[^32^P]-labelled RNA fragment and compared it to an unmodified reference **1** (Figure 2B). ASO/RNA duplexes showed initial RNA hydrolysis within 5 min releasing shorter 5’-[^32^P]-RNA products. Complete degradation of the substrate was observed after 30 min for unmodified **1**, while only 50% of the substrate was degraded after 60 min. The densitometric analysis revealed a two and a half times longer hydrolysis of RNA in the duplex with B-ASO^LNA^ (**9**) (t_1/2_ = 20 min, Figure 2B) compared to **1** (t_1/2_ = 8 min, Figure 2B and S2B). To evaluate the thermodynamic stability of duplexes of different B-ASOs (**1**, **3**, **5**, **7**, **9** and **11**) with their complementary DNA and RNA counterparts (labeled **1** or **2**, depending on the ASO strand length, Fig. 1D), we performed temperature-dependent UV-monitored melting experiments. The melting temperature and Gibbs’ free energy (ΔG) data are shown in Fig. 2C and Fig. S2C, respectively. The results clearly indicate that incorporation of FESANs into the ASO chain only slightly reduces the stability of the duplexes (as can be seen for the duplexes with B-ASO (**3**) compared to the duplexes with **1**). The inclusion of a cholesterol moiety in the oligonucleotide strand reduces the stability of the corresponding duplexes of **5** or **7** with DNA and RNA complements. The duplexes with LNA modification **9** and **11** are three base pairs longer than their parental models and therefore their thermodynamic stability is increased by the two components, the longer duplex length and the rigid C3’-*endo* LNA conformation.^18^ The Tm values of the B-ASO^LNA^ (**9**) and B-ASO^LNA^-CHOL (**11**) duplexes with RNA-2 increased to 81°C and 72°C, respectively. The presence of LNA flanks in the duplex of **11** to some extent compensates for the negative influence of the CHOL molecule on the thermodynamic stability of ASO/RNA (Figure 2C). Therefore, an appropriate combination of the ASO length (25-nt) as well as chemical modifications introduced into the sequence both contributed to the improved target affinity for B-ASO^LNA^-CHOL (**11)**.

**Figure 2.**
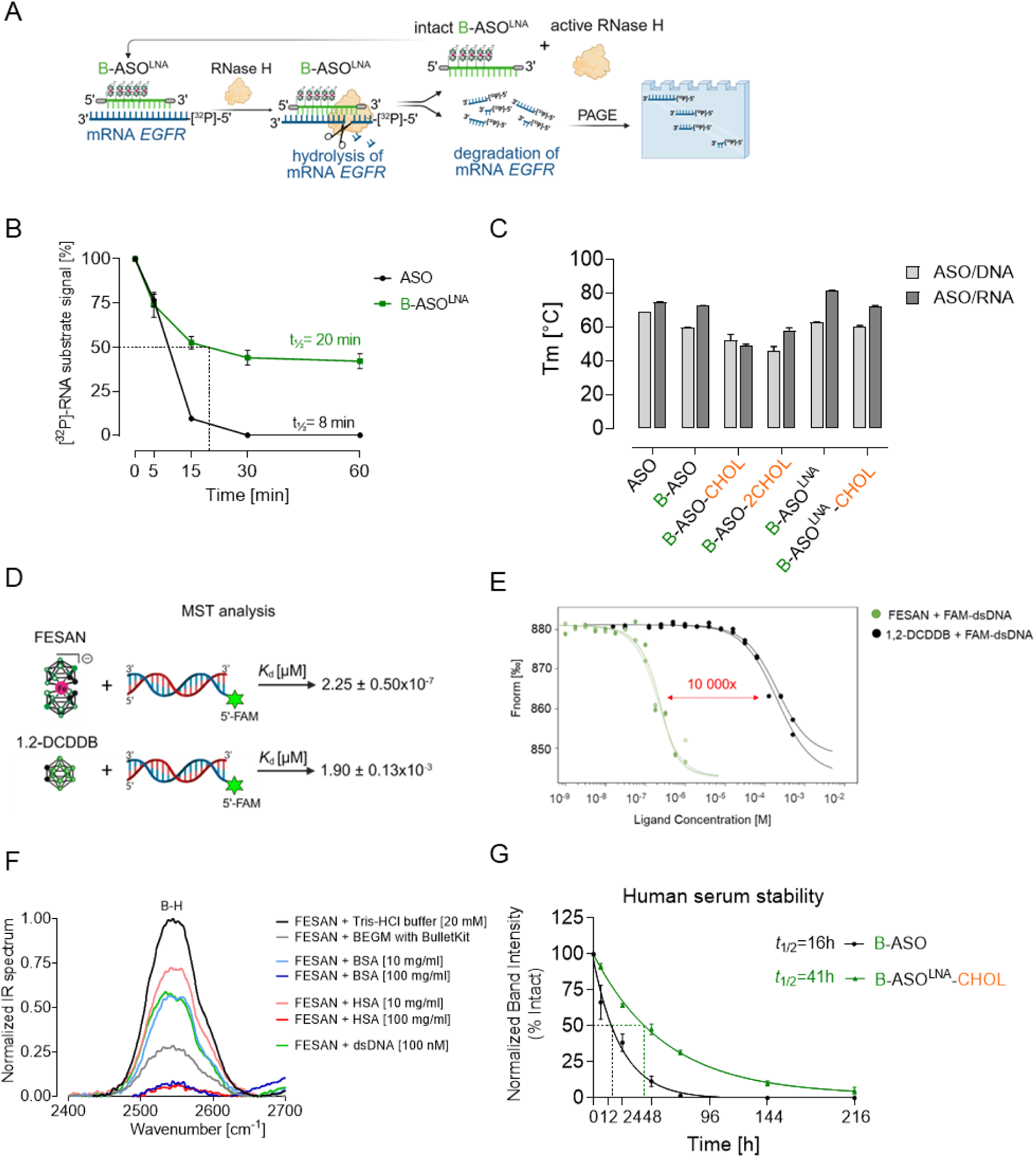
Biophysical and biochemical properties of boron cluster (FESAN) and B-ASO^LNA^-CHOL. (**A**) Illustration of the mechanism of the RNase H-mediated mRNA cleavage within B-ASO^LNA^/RNA duplex; (**B**) Time course of the RNase H-mediated endonucleolytic activity in the presence of B-ASO^LNA^/5′-[^32^P]-RNA heteroduplex analyzed by denaturating PAGE; (**C**) Melting temperature values (Tm) for various compounds tested (**1**, **3**, **5**, **7**, **9**, **11**) in the form of duplexes with complementary DNA or RNA; (**D**, **E**) Schematic representation of the affinities of two types of boron clusters, uncharged 1,2-DCDDB (0) and charged FESAN (−1) to short double-stranded DNA with fluorescent dye (5’-FAM) analyzed by MST; (**F**) IR spectra of FESAN in solutions containing a representative mixture of biomolecules with bovine serum albumin (BSA), human serum albumin (HSA), or bronchial epithelial cell growth medium (BEGM), as well as a control (FESAN in Tris-HCl buffer); (**G**) Stability of the chemically modified B-ASO^LNA^-CHOL (**11**) and unmodified B-ASO (**3**) was evaluated by incubating the samples in 50% human serum at 37°C for various times (ranging from 0 to 216 h), as analyzed using a 15% PAGE.

The therapeutic effect of BNCT requires efficient binding of ^10^B isotopes to the nuclear DNA of the cancer cells. Using miscroscale thermophoresis (MST), we found that FESAN ([Fe(C_2_B_9_H_11_)_2_]^-^) had an approximately 10^4^-fold stronger affinity for short double-stranded DNA labeled with a fluorescent dye (dsDNA^FAM^) compared to 1,2-dicarba-*closo*-dodecaborane (1,2-DCDDB, C_2_B_10_H_12_) at 25 °C (Kd = 2.25 ± 0.50×10^-7^ µM and Kd = 1.9 ± 0.13×10^-3^ µM, respectively) in ion buffer (Figure 2D and 2E).

Boron clusters exhibit unusual and robust B-H streching vibrations in the infrared (IR) spectroscopy in the range of 2450-2650 cm^-1^ which allow for compound detection (Figure 2F, black line).^19^ When tested in various biological solutions, the B-H peak of FESAN was detected in buffers containing representative biomolecules compared to the control. The B-H peak intensity decreased with increasing protein concentration. At 10 nM, BSA and HSA reduced the normalized B-H peak to approximately 60% and 70% of the control, respectively. FESAN combined with dsDNA also showed a B-H peak, but with a 40% reduction in intensity. This suppression likely results from the coordination of boron cluster anions and their metal-substituted derivatives with solvents, as well as organic or inorganic ligands, depending on the metal involved.^20^ Finally, the B-ASO^LNA^-CHOL (**11**) conjugate demonstrated a significantly prolonged half-life of almost 2 days when incubated with human serum (T₁/₂ = 41 h), in contrast to the unmodified B-ASO (**3**), which exhibited a half-life of only 16 h (Figure 2G).

### Crystal structure of human serum albumin-FESAN complex

To investigate the interaction between FESAN and human serum albumin (HSA), we determined the structure of the complex by X-ray crystallography. Co-crystallization of FESAN and defatted HSA in a molar ratio of 5:1 (compound:protein) resulted in crystals in space group I2 with different dimensions of the unit cell than in the Protein Data Bank. The asymmetric unit contained a single HSA molecule and provided the structure of the HSA-FESAN complex. Fourier difference electron density maps generated after molecular replacement phasing clearly showed an unoccupied electron density fragment within the IB subdomain that corresponded to the binding site of a FESAN molecule (Figure 3A). Additional ligands bound to the HSA molecule were identified, including residual fatty acids and compounds used for crystallization (sulfate ions, DMSO) and cryoprotection (glycerol). Refinement of the complex using data with a high resolution limit of 2.6 Å resulted in a well-validated model with favorable stereochemistry and R-factor and Rfree values of 0.234 and 0.314, respectively. Details on data collection and refinement are shown in Table S3A and B.

**Figure 3.**
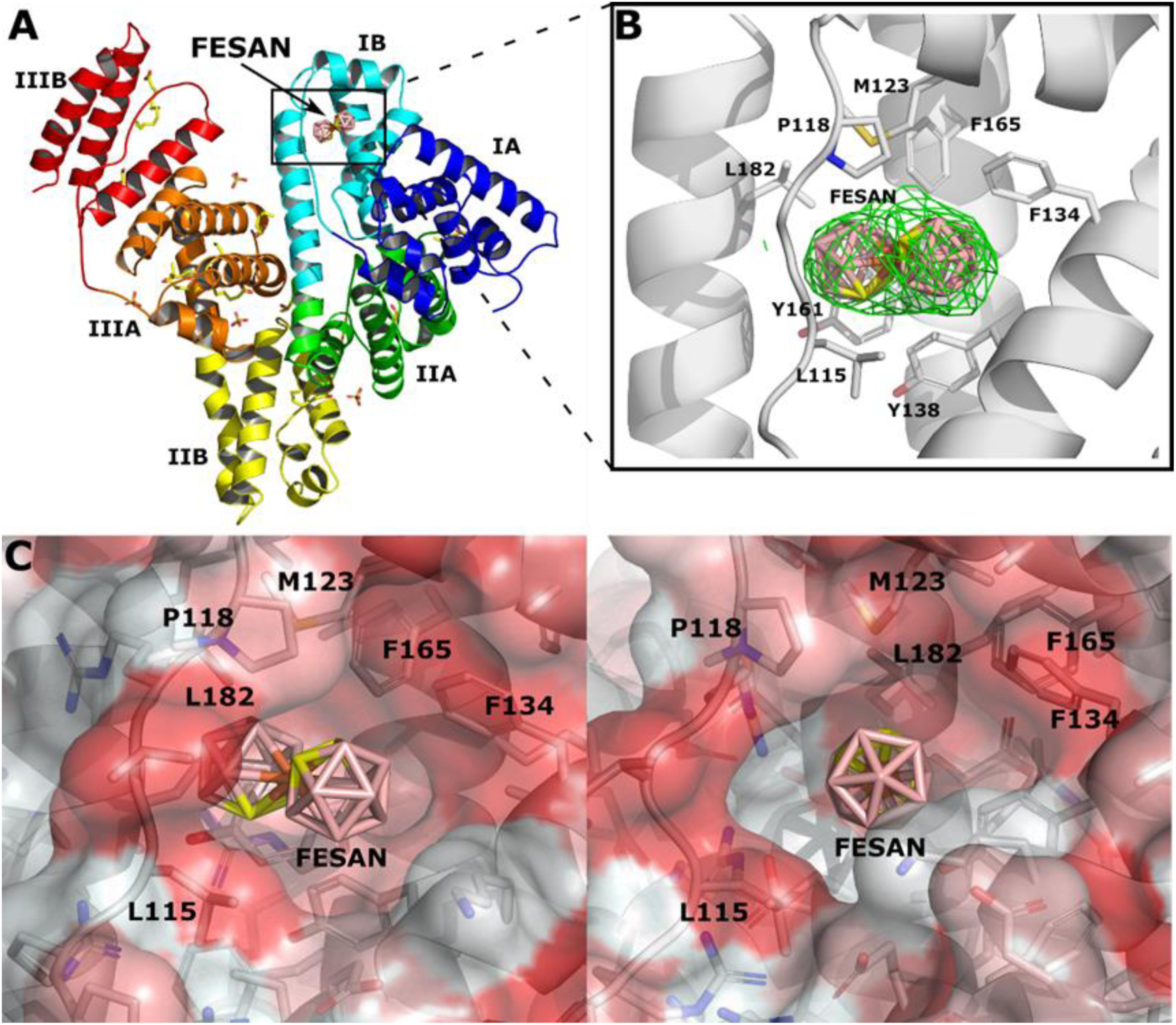
Crystal structure of human serum albumin (HSA) in complexes with FESAN. (**A**) The boron cage (FESAN) is shown in sticks and is colored in salmon. The remaining bound molecules are shown in sticks and are colored by atom type (C, yellow; O, red; S, light orange). (**B**) *F*O-*F*C omit map contoured at 3.0σ showing FESAN bound to the IB subdomain of HSA. FESAN is shown in a stick representation with atoms colored by atom-type: B, pink; C, yellow; Fe, orange. The subdomain IB is shown as a ribbon (light grey) with selected amino acids of the binding pocket shown in stick representation. The overall structure of HSA-FESAN with the subdomains rendered in different colors (domain IA, in blue; domain IB, in cyan; domain IIA, in green; domain IIB, in yellow; domain IIIA, in orange; domain IIIB, in red). (**C**) View of the fit of FESAN to the contours of the binding pocket in IB subdomain (shown as a salmon-colored, semi-transparent surface colored according to the hydrophobicity of the HSA chain). The view rotated by ∼45° around a vertical axis is shown on the right.

Our investigation of the structure of the FESAN-HSA complex^21^ revealed that the ligand is bound to site 3 within the IB subdomain. Binding site 3, which is enclosed by four alpha helices and covered by an elongated loop, consists of a combination of hydrophobic and polar residues that facilitate the recognition of saturated and unsaturated fatty acids,^22,23,24^ heme,^25^ bilirubin,^26^ anticancer drugs,^27,28,29^ and other drugs.^30,31^ The boron cage (FESAN) was predominantly located in the hydrophobic pocket formed by residues Leu115, Pro118, Met123, Phe134, Phe165, and Leu182 due to its lipophilic properties (Figure 3B and 3C). Interestingly, the presence of FESAN led to a conformational change of Tyr161, which adopted an “unstacking” state, while the position of Tyr138 remained unchanged in the “stacking” conformation (Figure S3C-E). This characteristic behavior was not observed in any of the previously studied complex structures in which ligands were bound in the IB subdomain (Figure S3C-E).

### B-ASO^LNA^-CHOL promotes cancer cell-specific targeting and uptake

In line with our hypothesis, EGFR^high^ human cancers were treated with the B-ASO compound to assess the ability to saturate with boron atoms by inductively coupled plasma mass spectrometry (ICP MS) (Figure 4A). The data showed that B-ASO applied in 0.5, 1 and 2 nM concentrations was rapidly taken up by EGFR^high^ cancer cells, especially after a 1 h starvation period followed by a 12 h treatment (Figure 4B). The highest boron concentration was observed in human skin (A431) and liver cancer cells (HepG2), which reached B: 33 µg/g (^10^B: 6.3 ug/g) and B: 26 µg/g (^10^B: 5 µg/g), respectively (Figure 4B). Conversely, cells that were not starved prior to treatment (0 h) or underwent a 6 h starvation had lower boron concentrations (Figure S4A).

**Figure 4.**
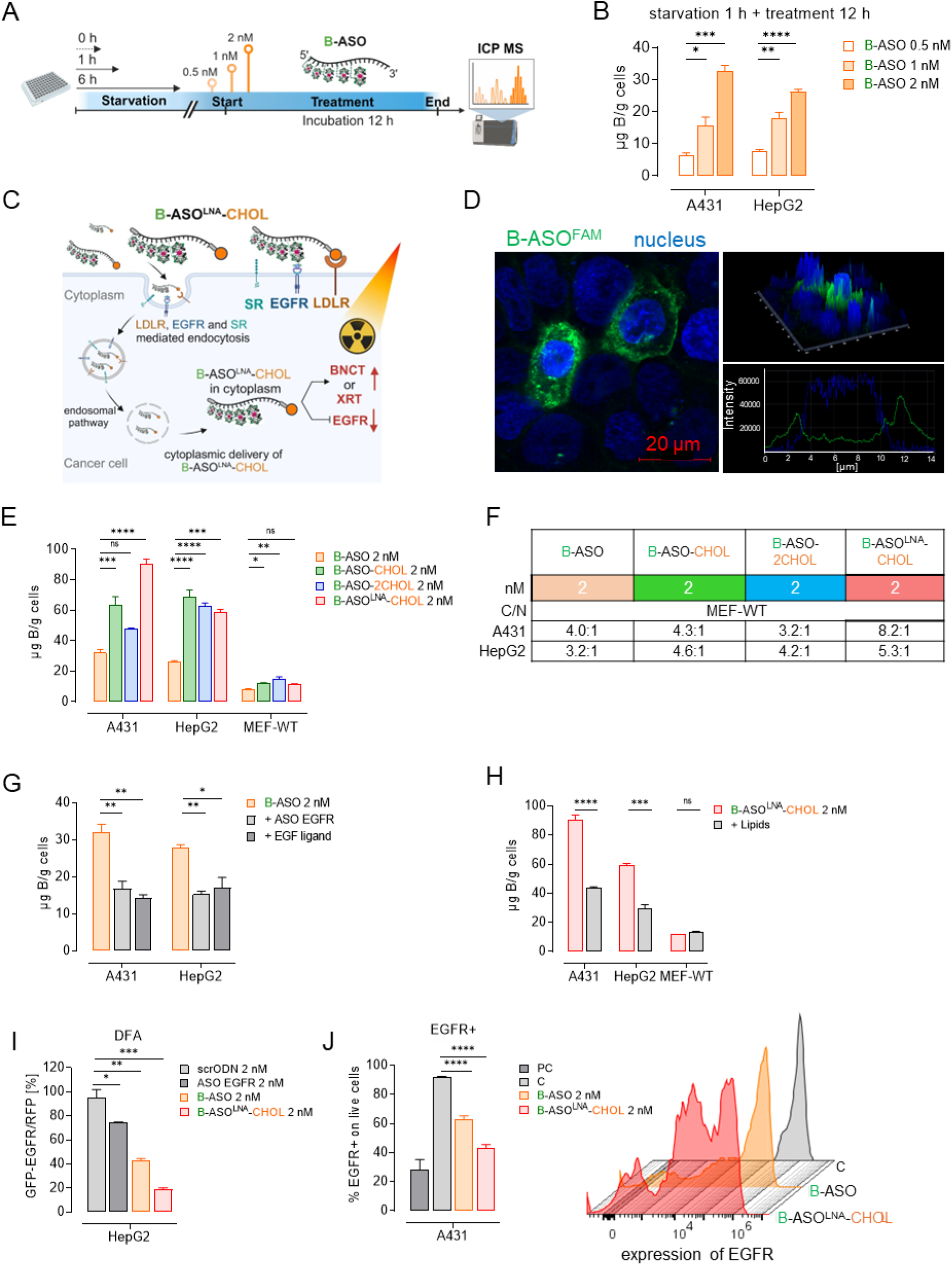
B-ASO^LNA^-CHOL effectively inhibited the EGFR expression while saturating human cancer cells with boron atoms. (**A**) Schematic representation of short-term starvation (STS, 1 h) and treatment (12 h) of cancer cells with B-ASOs analyzed by ICP MS; (**B**) Intracellular boron concentration in A431 and HepG2 after (STS 1 h + 12 h) with B-ASO in 0.5-2 nM concentrations; (**C**) Illustration of the uptake of B-ASO^LNA^-CHOL (**11**) via receptor mediated pathways by EGFR, LDLR or SRs in cancer cells; (**D**) The intracellular localization of B-ASO^FAM^ (**12**, green) and the nucleus (DAPI, blue) examined by confocal microscopy (magnification x80) after 1 h of incubation on A431 cells; Distribution shown by the topography scan, visualized with a scale of 20 mm; (**E**) Boron concentration in A431 and HepG2 and normal MEF-WT cells after incubation (STS 1 h + 12 h) with (**3**, **5**, **7**, **11**) (2 nM) verified by ICP MS compared to untreated control cells; (**F**) Therapeutic ratios for treatment with various of B-ASO conjugates between cancer and normal (C/N) cells after incubation (STS 1 h + 12 h); (**G**) Cellular uptake of B-ASO (**3**) measured by boron content, and EGFR activity was modulated either by reducing EGFR mRNA using ASO (**1**, 100 nM, 36 h, with Lipofectamine 2000) or by blocking the EGFR extracellular domain with EGF (10 ng/mL, 1 h); (**H**) Comparison of the uptake of B-ASO^LNA^-CHOL (**11**) (2 nM) via the LDLR in basic culture medium (30 µg/mL cholesterol and 3 µg/mL oleic acid) vs medium enriched with higher lipid concentrations (150 µg/mL cholesterol and 15 µg/mL oleic acid); (**I**) *EGFR* gene expression levels in HepG2 cells treated with (**1**, **3**, **11**) or scrODN control (**13**) (2 nM) determined by dual fluorescence assay (DFA) after 48 hours. Relative EGFP-EGFR to RFP fluorescence in cells transfected with plasmids alone set as 100% (C-control); (**J**) EGFR expression on A431 cells after treatment with B-ASO (**3**), B-ASO^LNA^-CHOL (**11**) (2 nM) and positive control (PC, PD153035-EGFR inhibitor, 1 µM) at 48 h, confirmed by flow cytometry analysis.

A431 and HepG2 cells exhibit significant expression of receptors on their cell membranes, such as EGFR, LDLR or scavenger receptors (SRs) (Figure 4C).^16,32,33^ Confocal microscopy showed that B-ASO^FAM^ (**12**) was localized in the cytosol of A431 cells within 1 h (Figure 4D and Figure S4B). Remarkably, A431 and HepG2 cells demonstrated effective uptake of B-ASO-CHOL (**5**) and B-ASO^LNA^-CHOL (**11**) (2 nM), resulting in boron concentrations of B: 64 to 92 µg/g (^10^B: 12.1 to 17.5 µg/g) and B: 67 to 100 µg/g (^10^B: 12.7 to 19.0 µg/g), respectively (Figure 4E), compared with other cancers (Figure S4C). The incorporation of double-CHOL groups (B-ASO-2CHOL, **7**) into the oligonucleotide chain did not improve delivery compared to the single-modified B-ASO-CHOL (**5**) (Figure 4E and Figure S4C). These compounds showed significant selectivity differentiating between cancer and normal (C/N) cells, with ratios ranging from 2.4:1 to 8.2:1 (Figure 4F). Other cell lines, including human cervical (HeLa) and breast cancer cells (MCF-7), exhibited increased uptake of B-ASO-2CHOL (**7**) at ^10^B: 6.5 µg/g (Figure S4D), thus their efficiency for BNCT remains insufficient. In contrast, the tested compounds showed low uptake in normal cell lines such as MEF-WT (mouse embryonic fibroblasts) and CCD-841 (human colon epithelial cells), which naturally have EGFR^low^ expression (Figure 4E and Figure S4C, S4D).^16^ Longer incubation times and higher concentrations resulted in a saturation of cancer cells with B-ASO inhibitors (Figure S4E-G). We investigated whether the altered expression of EGFR in A431 and HepG2 cells affected the uptake of B-ASO (**3**) using two methods: by inhibiting EGFR expression or by blocking the EGFR external domain. The results showed significantly lower B-ASO uptake when using the ASO EGFR inhibitor (B: ∼17 µg/g) or the EGF-free ligand (B: ∼16 µg/g) compared to cells treated with B-ASO (**3**) as a control (B: ∼30 µg/g) (Figure 4G). We also investigated whether the cholesterol moiety in B-ASO^LNA^-CHOL (**11**) enhanced the effect by interacting with the LDLR. The results suggested lower efficacy of boron atom delivery in a lipid-rich medium, with A431 cells being more sensitive than HepG2 cells (Figure 4H).

Overall, B-ASO^LNA^-CHOL (**11**) demonstrated significant inhibition of EGFR expression in a DFA assay performed on HepG2 cells, achieving about 80% inhibition compared to the scrODN control (Figure 4I). To further verify the activity of B-ASO^LNA^-CHOL agent (**11**), flow cytometry was also performed, which confirmed its EGFR-inhibitory effect on A431 cells (Figure 4J).

### B-ASO^LNA^-CHOL combined with BNCT enhances cytotoxicity in cancer cells

We assessed whether B-ASO^LNA^-CHOL–mediated BNCT enhances the therapeutic efficacy using thermal neutron beam irradiation (IRR), thereby overcoming the limited long-term effectiveness of monotherapy. B-ASO^LNA^-CHOL (**11**) inhibitor approach ensures free uptake of boron agents while sensitizing and weakening cancer cells by targeting EGFR mRNA prior to radiation. In our studies, we incubated A431, HepG2 and normal fibroblasts (BALB 3T3) with a single dose of B-ASO^LNA^-CHOL (**11)** or B-ASO (**3)** (2-8 nM concentration), followed by IRR of the cells with a 30-kW neutron beam for 15 min (Figure 5A). Prior to this, the survival rate of the tested cells at 7 days after IRR (without treatment), as determined by the clonogenic assay, was unaffected by IRR for up to 15 min (Figure 5B). Next, we tested whether A431 and HepG2 cells responded to treatment with B-ASO (**3**) (Figure 5C, E) and B-ASO^LNA^-CHOL (**11**) (Figure 5D, F) when exposed to IRR. We observed a significant decrease in the survival rate of the cancer cells treated with B-ASO^LNA^-CHOL (**11**) and IRR (Figure 5D, F), even at the lowest concentration of 2 nM oligonucleotide compared to controls treated with **3** (Figure 5C, E). The combination of B-ASO^LNA^-CHOL with BNCT showed an enhanced cytotoxic effect, leading to the elimination of both cancer cells. A comparable killing effect was observed in cancer cells treated with B-ASO (**3**) at the fourfold higher concentration (8 nM) after IRR (Figure 5C, E). Furthermore, we confirmed that BALB 3T3 exhibited high survival rates when treated with B-ASO^LNA^-CHOL (**11**) or B-ASO (**3**), with or without BNCT (Figure 5G and 5H). Due to the high affinity of FESAN for dsDNA, treatment with B-ASO^LNA^-CHOL (**11**) in combination with BNCT should exert DNA damage in A431 cells (Figure 5I). May-Grünwald Giemsa staining showed that the combination of B-ASO^LNA^-CHOL (**11**) with BNCT induced greater DNA fragmentation than treatment alone (Figure 5J), activating programmed cell death (Figure 5K). These results suggest that B-ASO^LNA^-CHOL (**11**) inhibitor may be a promising candidate for BNCT treatment.

**Figure 5.**
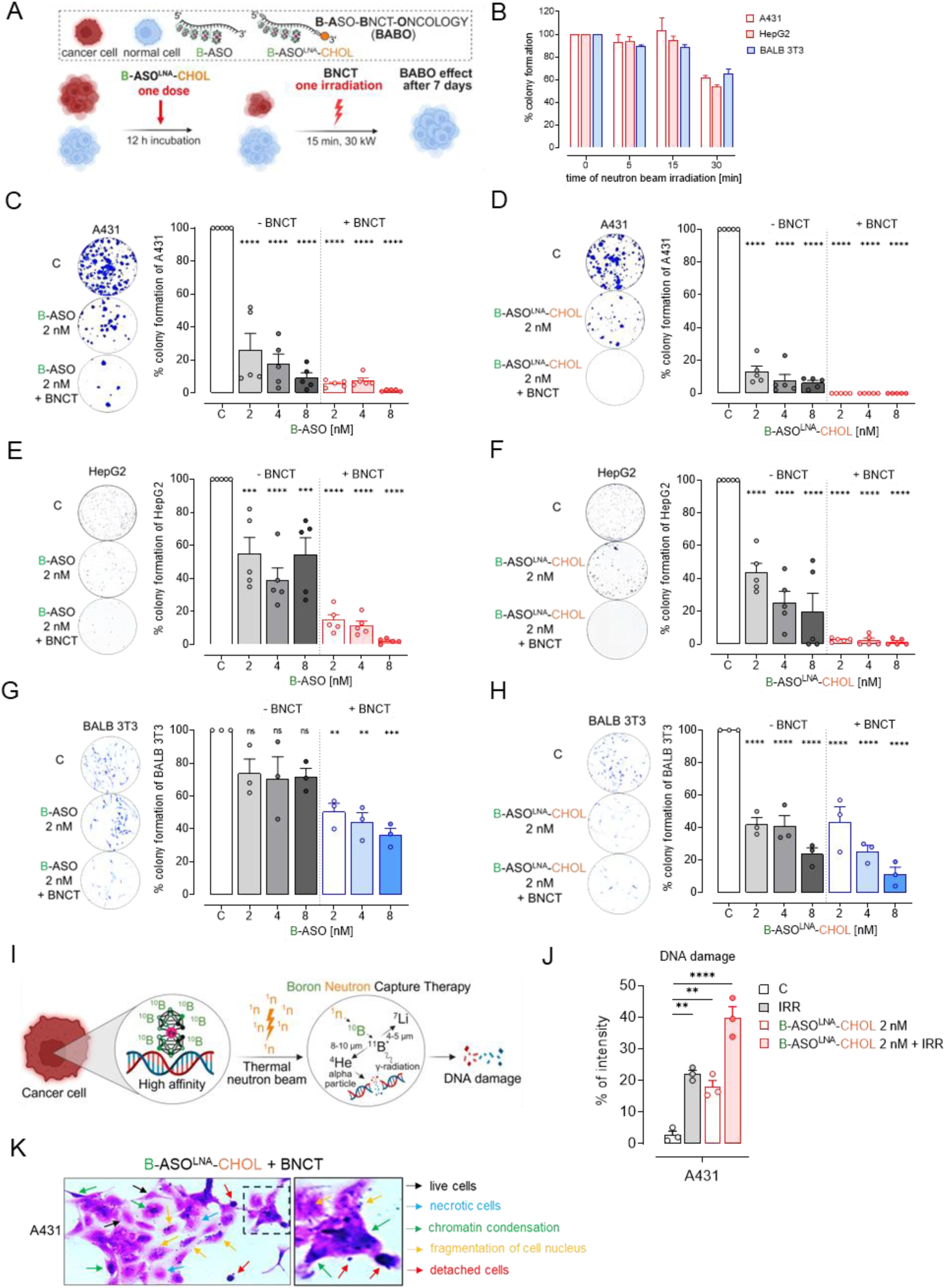
The combination of B-ASO^LNA^-CHOL with BNCT showed enhanced anticancer activity. (**A**) The design of dual B-ASO-BNCT-ONCOLOGY (BABO) therapy; (**B**) Survival of A431 and HepG2 cells and mouse fibroblasts (BALB-3T3) after thermal neutron irradiation of cells evaluated durations (0, 5, 15, 30 min at 30 kW) in the TRIGA MARK II Nuclear Reactor (Pavia, Italy); (**C-H**) Representative images of colony formation assays and survival graphs comparing the efficacy of B-ASO^LNA^-CHOL (**11**) and B-ASO (**3**) combinations with BNCT (15 min, 30 kW) in cancer and normal cells, C - untreated control cells; (**I**) Figure illustrating strong affinity of FESAN boron clusters on B-ASO^LNA^-CHOL (**11**) for nuclear DNA and impact on BNCT mechanism within a single cancer cell; (**J**, **K**) B-ASO^LNA^-CHOL (**11**) (2 nM) combined with BNCT increases DNA damage in A431 cancer cells, as shown by May-Grünwald-Giemsa staining after 72 h of incubation (scale bar, 40x magnification).

### EGFR-targeted radiosensitization by B-ASO^LNA^-CHOL conjugate in combination with XRT

Next, we investigated the radiosensitizing effect of the B-ASO^LNA^-CHOL (**11**) inhibitor in combination with XRT on human cancers overexpressing EGFR, including A431, HepG2, prostate (DU145), and head and neck squamous cell carcinoma cells (HNSCC_UM-SCC-1). Cells were treated with a single dose of B-ASO^LNA^-CHOL (**11**) together with a single dose of XRT (Figure 6A). The effect of these experiments was evaluated using the clonogenic assay in which cells viability was assessed 7 days after irradiation. The X-ray dose-dependent study showed that all cancer cells tested at a low dose of 0.25 Gy maintained 90% viability (Figure 6B). A higher radiation dose inhibited the colony formation of the cancer cells. EGFR levels in A431 cells treated with B-ASO^LNA^-CHOL (**11**) combined with XRT (0.25 Gy) were reduced by approximately 90% compared to the control (Figure 6C), as expected. In addition, the combination of FESAN (2 nM) with a low dose of XRT (0.25 Gy) slightly reduced EGFR expression in A431 cells after a 48 h incubation compared to boron cage treatment alone (Figure 6C). Finally, the results of the experiments B-ASO^LNA^-CHOL (**11**) or with FESAN at concentrations of 2, 4, or 8 nM, with or without XRT (0.25 Gy) indicated that the highly loaded B-ASO^LNA^-CHOL EGFR inhibitor effectively sensitized A431, HepG2, and DU-145 tumor cells to XRT (0.25 Gy) compared to FESAN monotherapy. This resulted in complete inhibition of colony formation in A431 cells treated with 2 nM B-ASO^LNA^-CHOL (**11**), and in HepG2 and DU145 cells treated with 8 nM, compared to control (Figure 6D-F). In radioresistant UM-SCC-1 cells, the combination of B-ASO^LNA^-CHOL (**11**) with XRT did not increase treatment sensitivity (Figure 6G). Our results suggest that the EGFR-targeted B-ASO^LNA^-CHOL inhibitor (**11**) effectively enhances cancer cell sensitivity to the cytotoxic effects of XRT.

**Figure 6.**
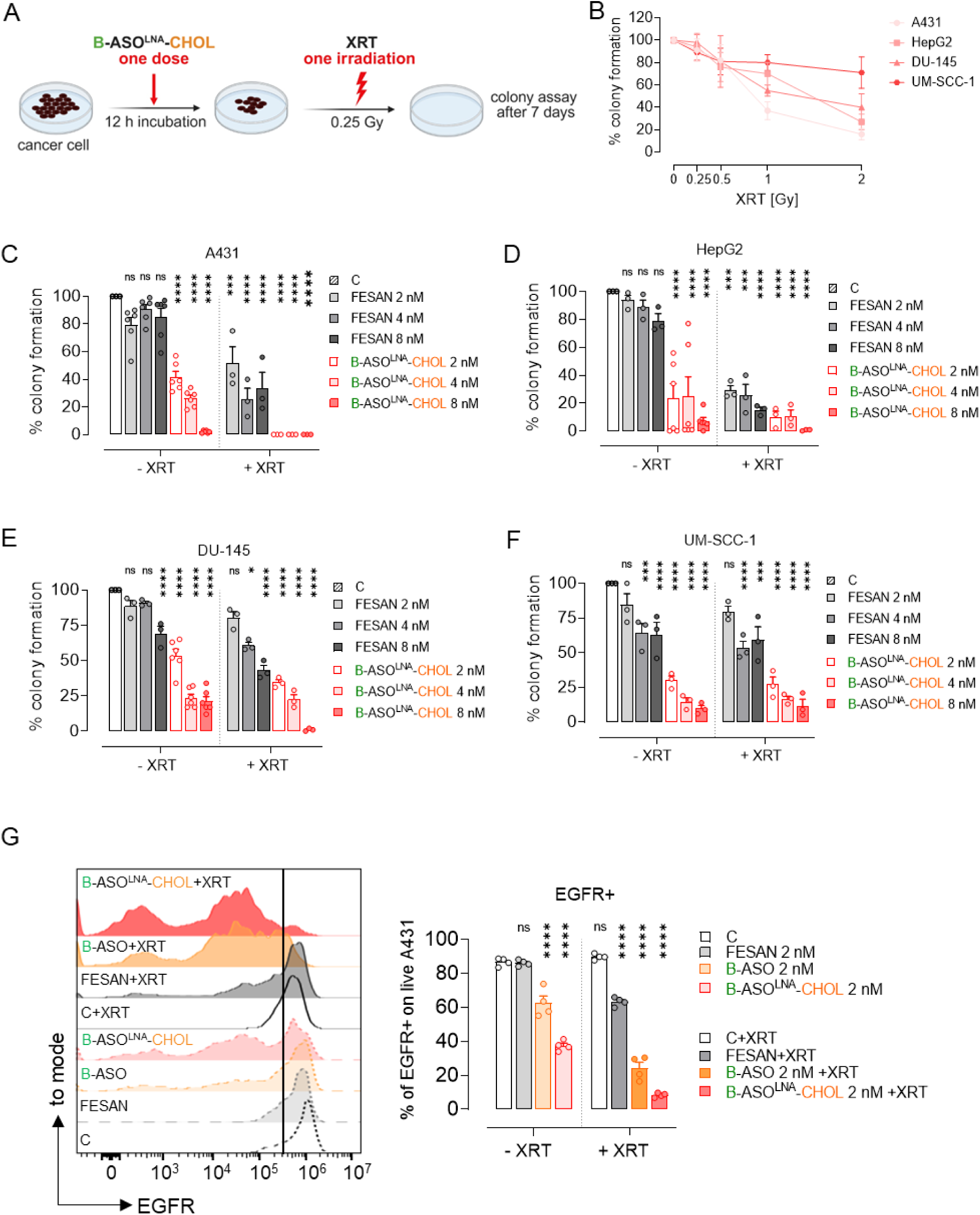
The combination of B-ASO^LNA^-CHOL and XRT sensitized human cancer cells to radiation. (**A**) The scheme illustrates the radiation sensitization effect of combining B-ASO^LNA^-CHOL (**11**) with XRT on EGFR^high^ human cancers; (**B**) The survival of A431, HepG2, DU-145, UM-SCC-1 cells evaluated after different doses of XRT (0, 0.25, 0.5, 1 and 2 Gy) using colony assay after 7 days. Experiments performed with the MultiRad160 irradiator (operating at 150 kV and 10 mA, SSD=37 cm with a temporal dose rate of 2.4 Gy/min); (**C**) EGFR expression in A431 cells confirmed by flow cytometry analysis after treatment with either B-ASO^LNA^-CHOL (**11**) or FESAN (2 nM) in combination with XRT (0.25 Gy) after 48 h. (**D-G**) The survival rate of tested cancer cells evaluated after 7 days using a colony assay to evaluate the efficacy of B-ASO^LNA^-CHOL inhibitor (**11**) or FESAN combinations with XRT (0.25 Gy), C - untreated control cells.

### B-ASO-CHOL induces macrophage activation acting as an immune adjuvant

The therapeutic efficacy of CpG ODNs, in combination with radiation, chemotherapy, or immune checkpoint inhibitors has made rapid progress in clinical development.^15^ The tested B-ASO-CHOL oligonucleotide (**5**) contains an unmodified class-B CpG ODN motif (Figure 7A).^34^ Therefore, we analyzed the surface expression of MHC-II, CD40 and CD80 as well as CD86 costimulatory molecules on mouse dendritic cells (DC2.4) (Figure 7B-E) and mouse macrophages (RAW 264.7) (Figure 7F) treated with B-ASO-CHOL (**5**) or FESAN using flow cytometry (Figure S5A).

**Figure 7.**
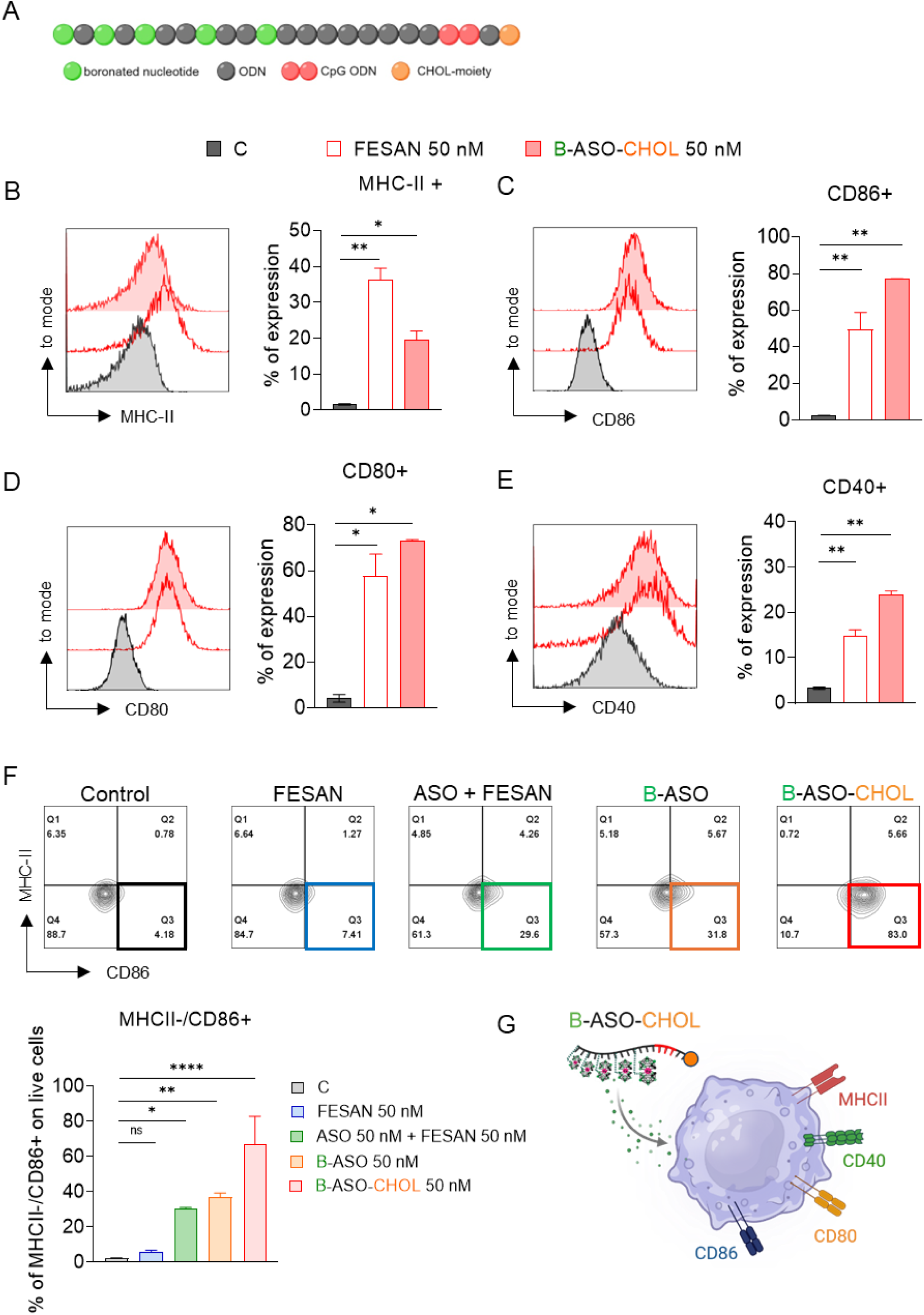
Enhanced expression of immunostimulatory molecules by B-ASO-CHOL was observed on dendritic cells and macrophages. (**A**) Schematic sequence of B-ASO-CHOL (**5**) with the immunostimulatory CpG ODN motif. Surface expression of MHC-II and the costimulatory molecules CD40, CD80, and CD86 on mouse dendritic cells (DC2.4 cells, **B-E**) and mouse macrophages (RAW 264.7 cells, **F**) analyzed by flow cytometry after treatment, C-untreated control cells; (**G**) Activated immune cells after treatment with B-ASO-CHOL (**5**).

B-ASO-CHOL (**5**) treatment resulted in upregulation of MHC class II, CD40, CD80 and CD86 molecules on DC2.4 cells after 24 h compared to CpG-1668 benchmark and other controls (LPS, DMXAA) (Figure 7B-E and Figure S5B). FESAN itself significantly enhanced the expression of key molecules on dendritic cells (Figure 7B-E). Next, we investigated whether a cocktail of unconjugated ASO (**1**) bearing an CpG motif with FESAN activates macrophages in vitro. The levels of the immunostimulatory molecule CD86+ induced by the cocktail (ASO (**1**) + FESAN) were comparable to those induced by the B-ASO (**3**) conjugate (Figure 7F). Finally, we confirmed that B-ASO-CHOL (**5**) enhanced the immunomodulatory effect by boosting MHCII-/CD86+ expression on RAW 264.7 macrophages compared to B-ASO (**3**) (Figure 7F). B-ASO-CHOL (**5**) slightly promoted the expression of MHCII+/CD86+ on RAW 264.7 cells (Figure 7F) providing opportunities for selective enhancement of immunotherapy, as demonstrated in Figure 7G. This finding highlights the need for further in-depth and targeted research on immunotherapy.

### Enhanced antitumor efficacy of B-ASO^LNA^-CHOL combined with local tumor radiation in vivo

Radiotherapy is a recognized alternative to surgery for patients with skin cancer who are not medically suitable for surgery, or for whom surgery could lead to a poor cosmetic outcome.^35^ Therefore, we investigated whether targeted inhibition of *EGFR* gene with B-ASO^LNA^-CHOL (**11**) enhances the cytotoxic effects of local XRT in vivo (Figure 8A). Immunodeficient mice (NSG) with established subcutaneously growing (s.c.) A431 human skin cancer were treated intratumorally with two doses of B-ASO^LNA^-CHOL (**11**) (0.5 mg/kg each) or PBS (control) with or without a single dose of local tumor irradiation (2 Gy) (Figure 8A and 8B). As shown in Figure 8C and 8D, injection of B-ASO^LNA^-CHOL (**11**) alone delayed but did not suppress A431 tumor growth, and the effect was similar to an XRT alone. However, the combination of XRT with targeted EGFR silenced by B-ASO^LNA^-CHOL (**11**) significantly enhanced A431 tumor regression in mice (Figure 8C, D and S6A). The harvested tumor (Figure 8E) and weight measurement (Figure 8F) after 18 days confirmed the efficacy of B-ASO^LNA^-CHOL+XRT (COMBO) therapy compared to monotherapy. Furthermore, the observed changes in tumor angiogenesis, as shown in Figure 8E, were confirmed by lower hemoglobin levels in the COMBO group compared to the control (PBS) (Figure 8G).

**Figure 8.**
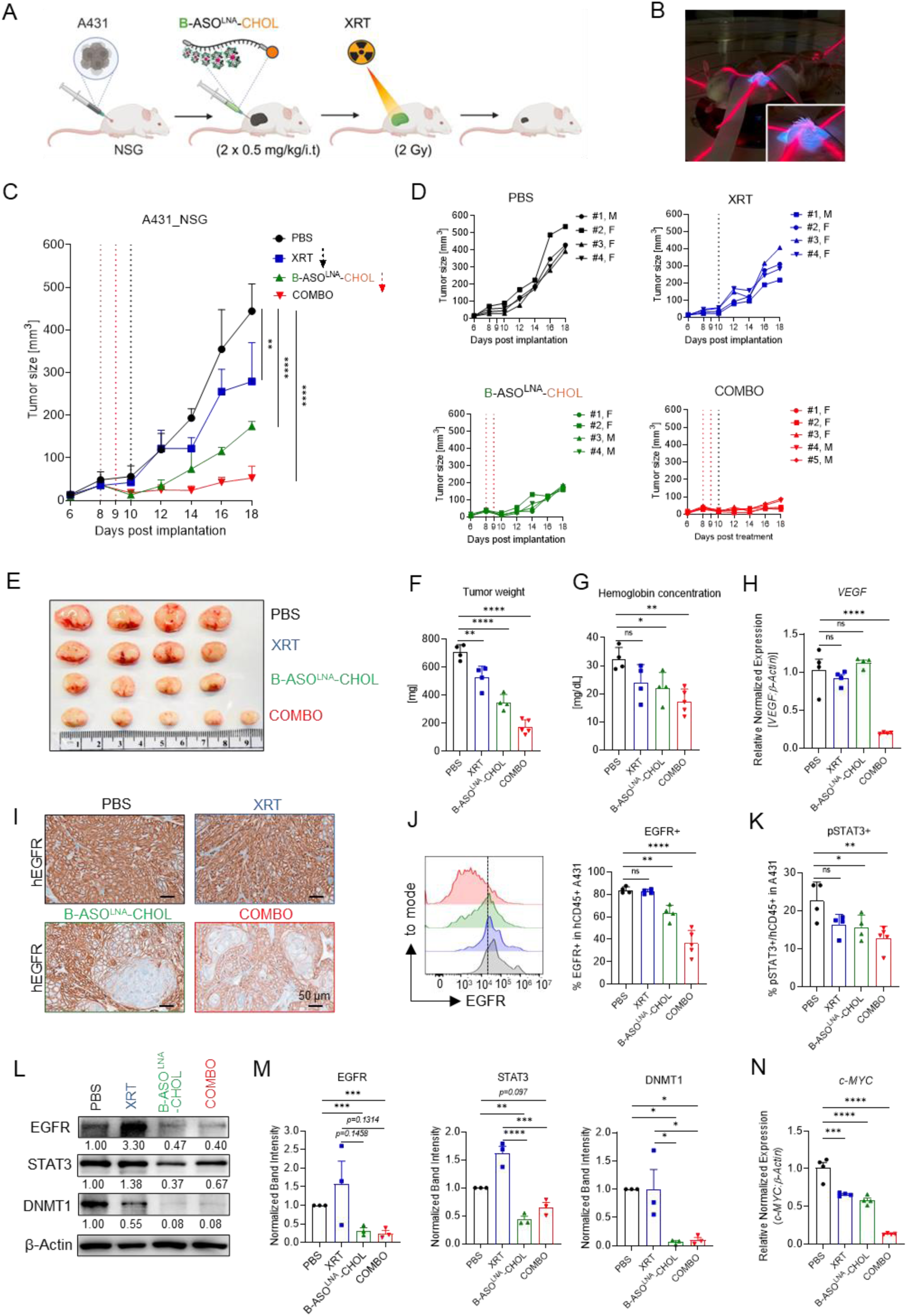
Tumor injections of B-ASO^LNA^-CHOL conjugate with local XRT effectively inhibited growth of xenotransplanted human skin tumor in immunodeficient mice. (**A**) Study design and treatment regimens for the efficacy studies; (**B**) local XRT with doses of 2 Gy using the MultiRad350 irradiator (operating at 350 kV and 10 mA, SSD=45cm with a temporal dose rate of 3.5 Gy/min) at the City of Hope Beckman Research Institute in Duarte, USA; (**C**, **D**) A431 cells implanted (s.c.) into immunodeficient mice (NSG) and treated after 8 days with two (i.t.) injections of B-ASO^LNA^-CHOL (**11**) (0.5 mg/kg) and/or an XRT dose of 2 Gy and PBS. The kinetics of tumor growth monitored using calipers (n=4-5 mice/group, M-male, F-female); (**E**) harvested tumor; (**F**) tumor weight at the end of the experiment; (**G**) tumor hemoglobin content indicating tumor vascularization measured by colorimetry; (**H**) VEGF expression measured by PCR; (**I**) representative histopathological sections of A431 xenograft stained after treatment with anti-human EGFR. Scale bar=50 μm; (**J**) expression of EGFR+ and (**K**) pSTAT3+ protein in hCD45+ A431 cells in tumor, measured by flow cytometry; (**L**, **M**) total protein levels of EGFR, STAT3, DNMT1, measured by Western blotting compared to β-actin (control); (**N**) c-MYC expression level measured by PCR.

Signal transducer and activator of transcription-3 (STAT3) is a downstream target of EGFR signalling. In turn, STAT3 promotes the expression of many tumorigenic genes including vascular endothelial growth factor (VEGF), a key angiogenic factor in cancer.^36^ COMBO successfully reduced *VEGF* levels in the tumor by approximately 80% compared to other groups (Figure 8H). Furthermore, a significant reduction in cell survival of approximately 80% was observed with the combined therapy compared to PBS measured by cytometry analysis (Figure S6B). To investigate whether COMBO affects the cellular expression of human EGFR in A431 xenografts, we performed immunohistochemical analysis of hEGFR and H&A in all treatment groups (Figure 8I and S6C). The results showed that hEGFR levels were significantly lower after treatment with the COMBO than with monotherapy (Figure 8I). A breast tumor was used as a control to validate the low EGFR expression compared to the A431 tumor (Figure S6C). Moreover, our evaluation showed that COMBO effectively reduced EGFR protein level in A431 xenograft by approximately 60%, as analyzed by both flow cytometry (Figure 8J) and western blotting (Figure 8L and 8M), similar to the effect observed with B-ASO^LNA^-CHOL (**11**) alone.

STAT3 is known to regulate expression of a key epigenetic regulator, DNMT1 in cancer cells.^37^ Our results showed that B-ASO^LNA^-CHOL (**11**) with or without XRT reduced pSTAT3/STAT3 and DMNT1 in tumor (Figure 8K-M). STAT1 and STAT3 generally play opposite roles in tumorigenesis, with STAT1 inhibiting tumorigenesis and being masked by STAT3 activity in cancer cells.^34,38^ The COMBO has shown a significant increase in antitumor STAT1 expression compared to other treatment groups (Figure S6D). Again, the increase of the *c-MYC* level is correlated with tumor invasion in EGFR^high^ patients in clinical trials.^39^ The combination of B-ASO^LNA^-CHOL (**11**) and XRT shows a synergistic effect, resulting in significant downregulation of *c-MYC* in A431 tumor cells compared to monotherapy (Figure 8N).

Overall, this study confirms that EGFR-targeted B-ASO^LNA^-CHOL (**11**) can enhance the inhibitory effect of radiotherapy against A431 xenograft in vivo.

## Discussion

Clinical trials of radiotherapy face obstacles, including the limited duration of tumor remission and the activation of resistance-promoting oncogenes. A new boron cluster-conjugated antisense oligonucleotide (B-ASO^LNA^-CHOL) was developed using click chemistry, a powerful tool of modern synthetic chemistry^40^, for use as an inhibitor of *EGFR* gene expression in combination with radiotherapy in cancer treatment.

The unique biological properties of metallacarborane (−1) (FESAN) allow them to penetrate biological membranes and localize effectively within the nuclei of cancer cells^41^, leading to a direct radiotherapy response.^5^ In addition, boron clusters are non-toxic and exhibit remarkable affinity for single- and double-stranded DNA^42^, with a binding strength approximately 10^4^-fold greater than that of uncharged boron clusters. This strong affinity is crucial for effective therapeutic applications, as highlighted by Viñas et al., demonstrated the importance of FESAN in facilitating initial electrostatic interactions with dsDNA through anti-coulombic “anion–anion” interactions.^43^

Moreover, boron clusters have the ability to bind plasma proteins, enhance their biodistribution and reduce renal filtration^44^, facilitating the delivery of B-ASO^LNA^-CHOL (**11**) to target tissues. Our results confirmed that the tested inhibitor has a strong affinity for serum albumin proteins, which is primarily due to hydrophobic forces between FESAN and proteins.^45^ Recently, E. N. Hoogenboezem et al., demonstrated that siRNA-lipid conjugates enhance albumin association, improving tumor bioavailability and facilitating extrahepatic and kidney inhibitor delivery in a breast xenograft.^46^

As far as we are aware, this study is the first to reveal the crystallographic structure of the FESAN-HSA complex^21^, demonstrating the strategic localization of the lipophilic boron cluster within a hydrophobic pocket defined by key amino acid residues. Additionally, our previous work indicated that FESAN can bind to hydrophobic amino acids within the binding domain of RNase H^14^, enhancing molecular interaction and enzymatic activity.^47^

B-ASO^LNA^-CHOL (**11**) has exhibited greater nuclease resistance than unmodified oligonucleotide, can improve a half-life profile and effective silencing of target gene in vivo.^48^ The incorporation of two ligands, FESAN and a cholesterol moiety, into B-ASO^LNA^-CHOL (**11**) significantly enhanced cancer cell-specific targeting and uptake. However, contrary to our expectations, the double cholesterol modification did not improve cellular uptake, possibly due to interactions between the hydrophobic FESAN and the cholesterol moieties positioned closely in the chain. Our findings indicated that the combination of B-ASO^LNA^-CHOL (**11**) with BNCT effectively inhibited the growth of A431 and HepG2 cancer cells at lower doses compared to unmodified B-ASO. Importantly, normal cells exhibited high survival rates both with and without BNCT, underscoring the therapeutic potential of this approach. As assumed, B-ASO^LNA^-CHOL (**11**), when combined with thermal neutron irradiation, significantly increased DNA damage, reflecting the effectiveness of BNCT.^1^

B-ASO^LNA^-CHOL (**11**) uptake appears to be dependent on EGFR density, as demonstrated by the greater sensitivity of A431 cells to the inhibitors compared to HepG2 cells. In related studies, strategies targeting EGFR^high^ glioblastoma cells have demonstrated similar efficacy while preserving the viability of normal fibroblasts when tested with an aptamer-boron cluster conjugate (*closo*-dodecaborate-GL44) in BNCT.^49^ Additionally, a study by Diego et al. underscores that boron cluster-loaded liposomes can enhance BNCT efficacy in ovarian cancer, indicating that boron-enriched compounds alone are insufficient for sustained therapeutic effect.^50^ Currently, Helsinki University Hospital in Finland^51^ and the Ibaraki Neutron Medical Research Center in Japan^52^ have initiated patient treatments using accelerator-based BNCT with the nuBeam® system.

Recently, Viñas et al. suggested that FESAN combined with XRT can inhibit cancer growth through a radiosensitizing effect.^5^ We confirmed that the B-ASO^LNA^-CHOL (**11**) in combination with XRT effectively inhibited EGFR protein and reduced the survival of human cancers compared to FESAN monotherapy. Additionally, the ability of FESAN to bind to specific amino acids in EGFR tyrosine kinase, as shown in our previous work^16^, may contribute to the observed direct cytotoxic effect following the XRT combination, though this finding requires further investigation.

Cancer immunotherapy has the potential to enhance the immune response through CpG-TLR9 activation against cancers and inhibit the progression of metastasis, thereby improving the long-term outcomes of combined cancer therapies.^53^ Our results have shown that B-ASO-CHOL (**5**) containing a CpG ODN motif significantly increased the population of activated antigen-presenting MHC-II/CD86+ macrophages and enhanced the expression of MHC class II and co-stimulatory molecules on dendritic cells. Notably, adding a cholesterol residue next to the unmodified CpG-motif did not reduce its immunostimulatory activity. Moreover, the activation of endosomal TLR9 by CpG-ODN^54^ may facilitate the enhanced release of B-ASO-CHOL (**5**) into the cytoplasm, further contributing to therapeutic efficacy. Yaxin Shi et al. showed that a boron capsule loaded with an immunological adjuvant combined with BNCT induced a systemic antitumor immune response against melanoma in vivo, effectively inhibiting cancer growth and remodelling the tumor-immune microenvironment.^55^

Finally, administration of B-ASO^LNA^-CHOL (**11**) combined with localized XRT in human A431 xenografts in NSG mice resulted in a pronounced and significant inhibition of tumor growth, surpassing the effects observed with either treatment separately. In NSG mice, deficient in T, B, and NK cell activity, the combination treatment resulted only in a cytotoxic effect on the A431 tumor cells, without the immunomodulation benefits of B-ASO^LNA^-CHOL (**11**). Immunohistochemical and cytometric analyses confirmed the inhibition of EGFR target in the B-ASO^LNA^-CHOL (**11**) with XRT group, aligning with western blot and flow cytometry data. Notably, XRT monotherapy resulted in a three-fold increase in EGFR, corroborating findings from other studies.^56^ STAT3 is crucial for tumor cell proliferation and survival (via EGFR), angiogenesis (via VEGF) and invasion, thus providing a direct link between oncogenesis and immune evasion.^57^ B-ASO^LNA^-CHOL (**11**), both with and without XRT, effectively inhibited STAT3 and DNMT1 in A431 xenografts. Recently, Wang et al. described a bifunctional TLR9-targeted decoy oligodeoxynucleotide (CpG-STAT3d) that led to the inhibition of STAT3 and DNMT1 targets, affecting the epigenetic control of acute leukemia in vivo.^37^ VEGF plays a crucial role in the early stages of tumor development and contributes to poor clinical outcomes.^36^ The hemoglobin concentration results confirmed decreased tumor vascularization in the COMBO group, along with inhibited levels of angiogenesis factor. Hyperproliferation driven by the proto-oncogene *c-MYC*^58^ was significantly inhibited by the combination of B-ASO^LNA^-CHOL (**11**) and XRT, similar to the proposed boron cluster-nanostructures strategy targeting both *EGFR* and *c-MYC* genes.^59^

## Materials and Methods

### Synthesis of oligonucleotides

The synthesis of alkyne-functionalized DNA oligonucleotides (sequences in Figure 1D) was performed in-house according to a standard procedure using the phosphoramidite solid-phase method.^16^ The LCA CPG glass support was used to obtain the U_PR_-ASO (**2**) oligonucleotide. The introduction of the cholesterol moiety forced the change of the support to CHOL-TEG-CPG (U_PR_-ASO-CHOL **4**; U_PR_-ASO-2CHOL **6**), while in the case of the LNA moieties, the oligonucleotide synthesis had to be performed with a universal support (U_PR_-ASO^LNA^ **8**, U_PR_-ASO^LNA^-CHOL **10**). The fully protected oligomers were deprotected and cleaved from the solid support by two methods: by treatment with aqueous ammonia (1 mL, 30%), overnight at 55 °C (standard conditions for LCA CPG and CHOL-TEG-CPG) or with a 1 mL solution consisting of 36 µL of 10 mM LiCl solution in 964 µL of 30% ammonia (universal support, 12 h, 65 °C). For compounds in which this cholesterol fragment was present (**4**, **6**, **10**), purification consisted of two purifications on C18 SepPak cartridges (Waters) using aqueous acetonitrile solutions (from 10% to 80%, approx. 2 mL each). The fraction containing the pure desired product was then desalted again using C18 SepPak cartridges, with the 5’-DMT group removed by treatment with 2% TFA and elution of the desired products with 80% aqueous acetonitrile (approx. 2 mL). The collected samples were concentrated and analyzed by ESI-Q-TOF mass spectrometry and analytical RP-HPLC. Oligonucleotides containing a 2’-O-propargyluridine moiety (Figure 1A-D) were modified with alkyl azide of FESAN^17^ after synthesis by performing a copper-catalyzed azide-alkyne cycloaddition reaction as described in previous work^60^.

### Crystallization, X-ray data collection, processing, structure solution, refinement and analysis

Prior to crystallization experiments, human serum albumin (HSA) from Sigma-Aldrich (# A1653) was mixed with a boron cluster (FESAN) dissolved in 20 % (v/v) DMSO to obtain an HSA-FESAN complex solution with a concentration of 1.7 mM. Crystals were grown by vapor diffusion with hanging droplets consisting of 2 μL protein and 2 μL well solutions containing 1.2-1.6 mM (NH_4_)_2_SO_4_ and 1% (v/v) FHIP (1,1,1,3,3,3-hexafluoro-2-propanol). The crystals formed after 3-5 days of incubation at 20 °C and were cryoprotected with glycerol mixed with mother liquor to a concentration of 25 % (v/v) and then cooled directly in the N2 stream. Diffraction data were collected using the Rigaku XtaLAB Synergy-S diffractometer equipped with a HyPix-6000HE Hybrid Photon Counting (HPC) detector and a sealed Cu microfocus X-ray source, and a low temperature Oxford Cryostream 800 liquid nitrogen cooling system at 100 K. The data acquisition strategy was calculated using CrysAlis PRO to ensure the desired data redundancy and percent completeness. The data was processed, integrated and scaled using CrysAlis PRO and AIMLESS.^61^ Data acquisition and processing statistics are shown in Figure S3. Molecular substitution was performed with HSA coordinates (PDB ID: 2XVV)^62^ using MOLREP.^63^ Modeling and molecular visualization was performed in COOT.^64^ Ligand constraints were calculated using Grade2 from Grade Web Server^65^ and refinement was performed using REFMAC5.^66^ All refinement steps were monitored with Rcryst and Rfree values. The stereochemical quality of the resulting models was evaluated using the program MOLPROBITY^67^ and the validation tools implemented in COOT. The values of the mean temperature factors for the protein backbone and side chains, ligands and water molecules were calculated using the program BAVERAGE from the CCP4 suite.^68^ The refinement statistics of the described structure are listed in Figure S3. All figures were created with PyMOL v.2.5.0.

### Cell culture

The cell lines (HeLa, A431, MCF-7, U87-MG, DU-145, UM-SCC-1, DC2.4, RAW 264.7, HTC-116, HepG2, MEF-WT, CCD-841 CoN and BALB-3T3) were cultured according to the ATCC protocol. The culture media were supplemented with heat-inactivated fetal bovine serum (FBS), 100 U/mL penicillin and 100 µg/mL streptomycin (at 37 °C and 5% CO_2_). Reagents from Gibco (BRL, Paisley, NY, USA) and ATCC (Manassas, Virginia, USA) were used. Cell cultures were tested for mycoplasma contamination using the EZ-PCR Mycoplasma Detection Kit (BI, Cromwell, CT, USA) and were negative. Determination of boron content after treatment in the cells by ICP MS was measured as described in our previous work.^16^

### Colony formation assay

The treated cells were irradiated with thermal neutron beams (approximately 2×10^9^ n/cm^2^/s at 30 kW) in the thermal column of the TRIGA MARK II Nuclear Reactor (University of Pavia, Italy) or with X-rays with the MultiRad160 irradiation device at the City of Hope Beckman Research Institute in Duarte (California, USA). After treatment and prior to irradiation, the cells were washed twice with PBS, the medium was replaced with fresh medium, and 500 cells were seeded into 6-well plates and incubated at 37 °C with 5% CO_2_ for up to 7 days. Cells were then washed with PBS, fixed in 80% methanol for 10 min and stained with 0.01% crystal violet (Sigma-Aldrich, Saint Louis, Missouri) for 30 min. Colonies of more than 30 cells were counted using ImageJ software (version 1.54).

### Confocal microscopy

Imaging was performed using an LSM 880 with Airyscan confocal microscope (Zeiss, Oberkochen, Germany). For imaging studies, A431 cells were seeded on 6-wells and incubated with B-ASO^FAM^ (**12**) and DAPI and then fixed with 2 % paraformaldehyde (Fisher Scientific, NH, USA). Slides mounted in Vectashield HardSet medium (#H-1400, Vector Laboratories, Burlingame, CA, USA) were visualized with an LSM 510-Axiovert confocal inverted microscope (Zeiss) and analyzed with LSM ImageBrowser (version 4.2.0.121; Zeiss).

### Transcriptomic and protein assays

For qPCR, total RNA was extracted from cancer cells using the Maxwell RSC simplyRNA Cells System (AS1390; Promega, Madison, WI), then reverse transcribed into cDNAs using the iScript cDNA synthesis kit (Bio-Rad). QPCR was then performed with specific primers for VEGF, c-MYC and β-Actin as previously described^10,69^ with a CFX96 Real-Time PCR Detection System (Bio-Rad). Western blots were performed with specific antibodies against EGFR (EP38-Y, Abcam, Cambridge, UK), STAT1, STAT3 (#9172, #9132, Cell Signaling Technology, Danvers, MA) or β-actin-HRP (horseradish peroxidase) (Sigma-Aldrich) and described previously.^10,67^ Blots were imaged in a Bio-Rad ChemiDoc MP system using Enhanced Chemiluminescence (ECL; SuperSignal West Femto Maximum Sensitivity Substrate) and the resulting images were analyzed using the provided Bio-Rad Image Lab software.

### Flow cytometric analysis

RAW264.7 and DC2.4 cells were treated with FESAN, ASO (**1**), ASO+FESAN, B-ASO (**3**), B-ASO-CHOL (**5**), CpG1668 of 50 nM and LPS (1 µg), DMXAA (0.1 mg) as positive control, and then incubated for 24 h. Cell surface staining was performed with fluorochrome-labeled antibodies specific for CD40 (#12040183), CD80 (#104726), CD86 (#11086285) and MHC-II (#17532182) after LIVE/DEAD staining (#L34966). Reagents from Invitrogen, BioLegend and eBioscience (CA, USA) were used. Fluorescence data were analyzed on NovoCyte Quanteon (Agilent) flow cytometers using FlowJo v.10 software (TreeStar, Ashland, OR).

### Animal studies

The animal study was conducted in accordance with established institutional policies and approved protocols of the Institutional Animal Care and Use Committee (IACUC) at City of Hope (COH, Duarte, CA). NOD/SCID/IL-2RγKO (NSG) mice aged 6 to 8 weeks, originally obtained from The Jackson Laboratory (Bar Harbor, ME), were maintained at COH. For the efficacy studies with the A431 tumor implanted subcutaneously (s.c.), 6 × 10^6^ A431 cells were resuspended at a 1:1 ratio with Matrigel (#356231, Corning, Corning, NY, USA) and 1×PBS, and SC was injected into female/male NSG mice (n = 4-5). Eight days after injection of A431 cells, the mice developed solid tumor with a volume of approximately 50 ± 10 mm^3^. To investigate the efficacy of XRT (2 Gy) and B-ASO^LNA^-CHOL treatment, the mice were treated with B-ASO^LNA^-CHOL (**11**) (0.5 mg/kg, intratumor (i.t.), daily for two days, PBS control (same schedule) and XRT under xylazine/ketamine anesthesia or a combination of B-ASO^LNA^-CHOL (**11**) and XRT with oligonucleotide treatment starting two days prior to XRT. Suspension of cells from the tumor were prepared according to our previous work.^70^ The hemoglobin concentration in the tumor samples was measured using a hemoglobin assay kit (MAK115, Sigma Aldrich, St. Louis, USA) before the ACK buffer was used. Extracellular staining was performed with fluorochrome-labeled antibodies against hCD45 (#2432599), hEGFR (#352910), Annexin V (#A35110) or intracellular pSTAT3 protein (#3177617), LIVE/DEAD staining (#L34966). Fluorescence data were analyzed on NovoCyte Quanteon (Agilent) flow cytometers using FlowJo v.10 software (TreeStar, Ashland, OR). Western blot and PCR analysis were performed as described above. Specific antibodies against hEGFR and hematoxylin and eosin (H&E) were used for histochemical staining of tumor sections. Reagents were from Invitrogen, BioLegend and eBioscience (CA, USA) unless previously marked.

### Statistical Analysis

An unpaired Student t-test was used to calculate 2-tailed P-values to estimate statistical significance between 2 treatment groups. One-way analysis of variance and Bonferroni post-test were used to assess differences between multiple groups and in tumor growth kinetics, respectively. Statistically significant P-values were indicated in the figures compared to untreated or PBS groups). Data are presented as mean±SD (n=3-5); *P < 0.05, **P < 0.01, ***P < 0.001, ****P < 0.0001 by one-way ANOVA with Bonferroni’s correction post hoc test. Data were analyzed using Prism software version 9 (GraphPad Software).

## Acknowledgments

This work was supported in part by the NCI/NIH grant number R01CA284593 (to M.K.). We acknowledge dedication of staff members of the Analytical Cytometry, Light Microscopy/Digital Imaging, DNA/RNA Synthesis, Research Pathology and Animal Facility Core Laboratories supported by the NCI/NIH grant number P30CA033572. The content is solely the responsibility of the authors and does not necessarily represent the official views of the NIH.

This research received funding from the National Science Center, Poland, NCN ETIUDA 8 grant [2020/36/T/ST4/00485 to D.K.], NCN Miniatura grant [2021/05/X/ST4/01145 to J.S.], and Rector’s grant by Silesian University of Technology, Gliwice [04/010/RGJ22/1035 to A.J-K. This research was supported by a CMMS PAS Grants for Young Scientists [551-11 (to D.K.) and 551-12 (to K.E.-O.)] and by statutory funds from the Centre of Molecular and Macromolecular Studies, Polish Academy of Sciences (to B.N.). The experimental activities with neutron irradiation were supported by the Italian National Institute of Nuclear Physics (INFN) (to S.A.), project ENTER-BNCT (to N.P.). The Rigaku XtaLAB Synergy-S X-ray diffractometer system used for the results included in this publication was supported by funding from the EU Regional Operational Program for the Lodz Region, RPLD.01.01.00-10-0008/18 (to R.D.)

The authors thank Anna Maciaszek (CMMS PAS) for the synthesis of oligonucleotides, Ewelina Wielgus (CMMS PAS) for the mass spectrometric analysis, Patrycja Szczupak (CMMS PAS) for the purification of the HSA and Bartłomiej Kost (CMMS PAS) for the IR analysis. The authors are grateful to Drs. Marcin Lis (UJ), Monika Bugno-Poniewierska (UJ), and Rafał Jakubowski (CMMS PAS) for their support, suggestions, and/or critical reading.

## Contributions

B.N., M.K., N.P., and D.K. conceived and designed the study; A.J.-K., J.S., J.H., D.W., E.K., R.D., A.S., K.E.-O., K.K., and D.K. performed experiments; K.E.-O., and K.K. synthesized the oligonucleotides, M.K., B.N., K.E.-O., J.S., J.H., R.D., and D.K. reviewed and edited the manuscript; M.K., B.N., A.J.-K., R.D., M.L., N.P., S.A., J.S., K.E.-O., and D.K. funded and supervised the study, and D.K. wrote the paper.

## Supplementary

**Figure S1.**
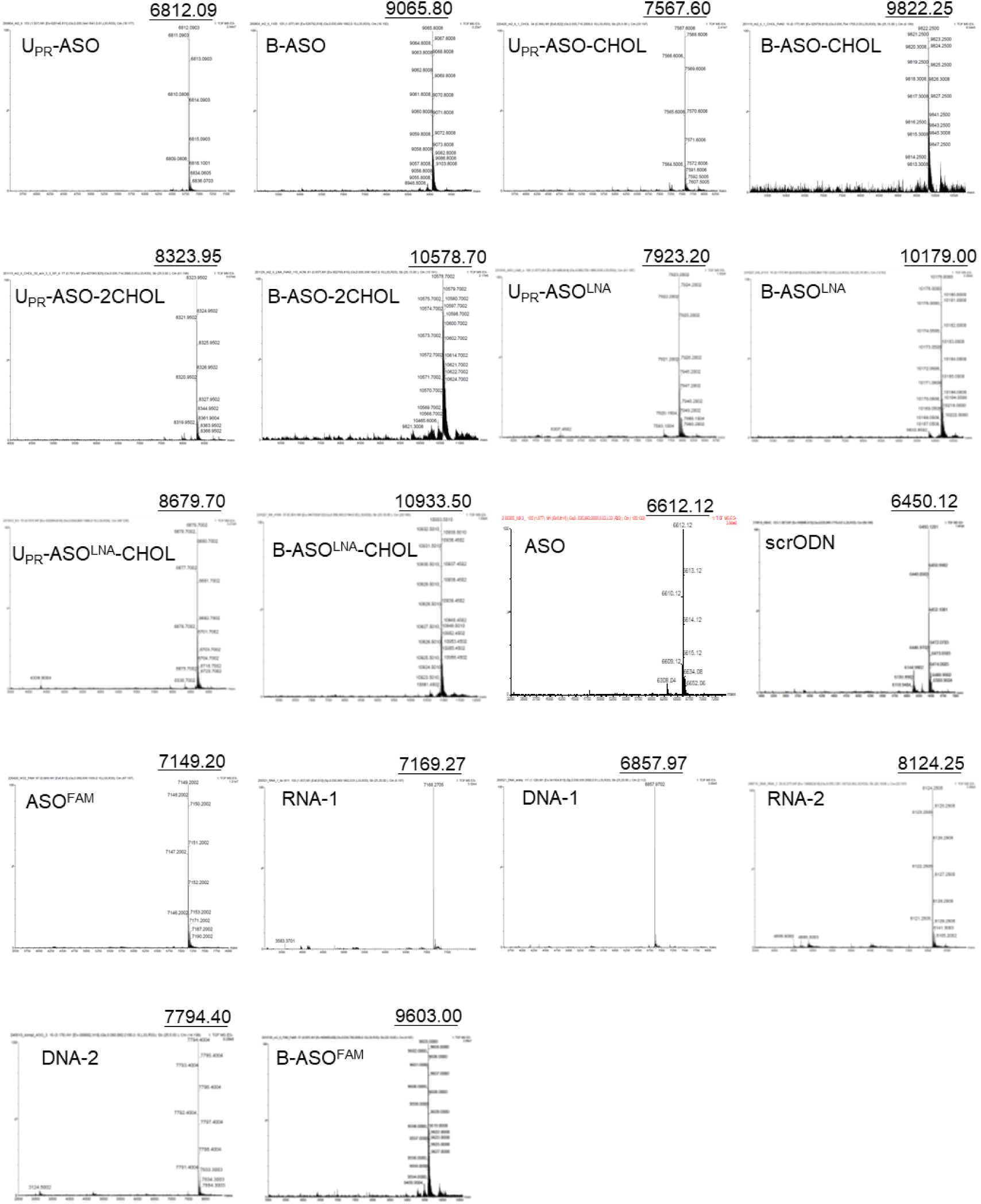
The sequence and purity of the obtained B-ASO^LNA^-CHOL conjugates and its derivatives were confirmed by analyzed by electrospray ionization quadrupole time-of-flight mass spectrometry (ESI-Q-TOF MS).

**Figure S2.**
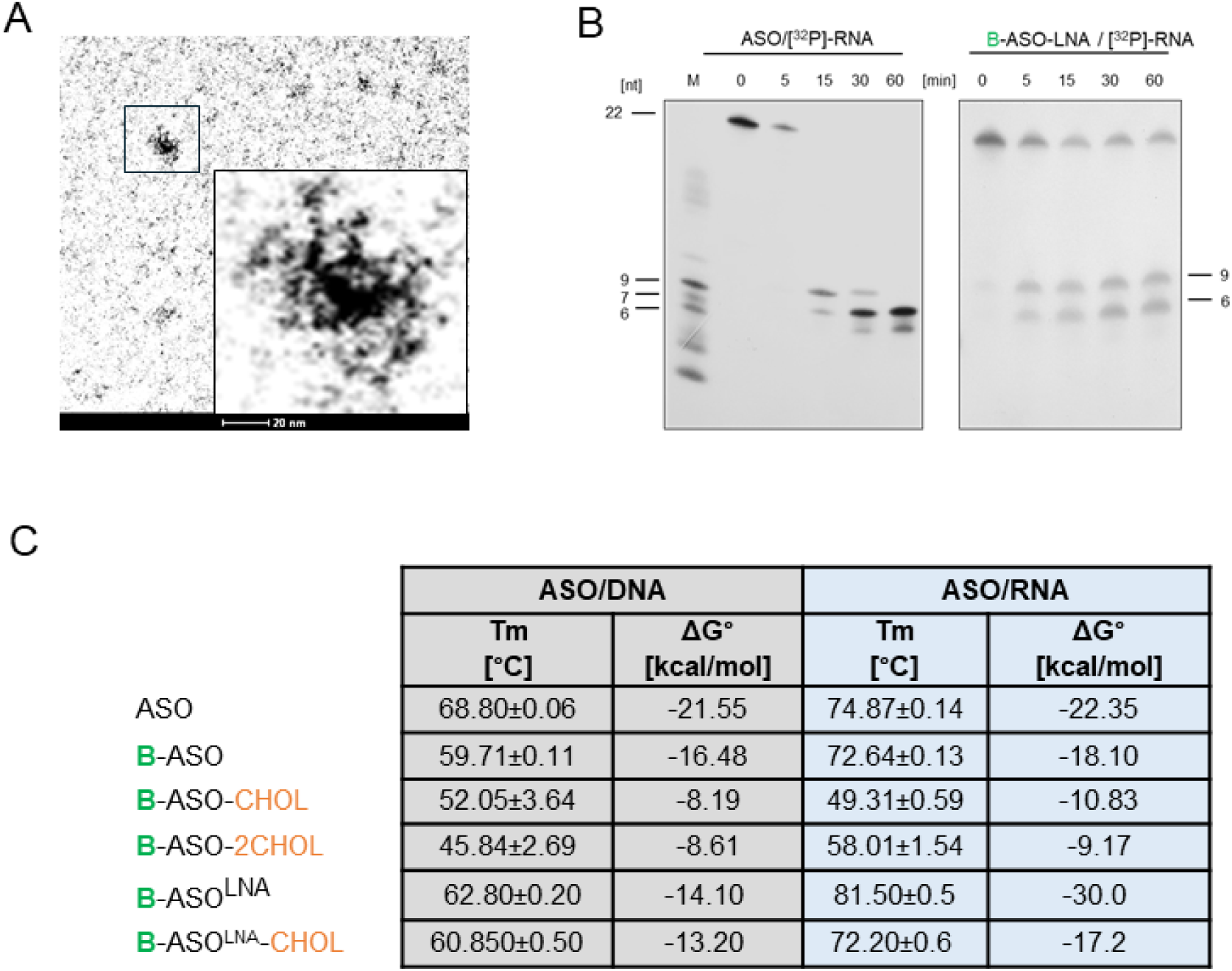
Characterization of physicochemical properties of B-ASO^LNA^-CHOL inhibitor. (**A**) Structural analysis of B-ASO was performed using Cryo-TEM; (**B**) PAGE analysis of RNase H-mediated hydrolysis of B-ASO^LNA^ and unmodified ASO with [^32^P]-RNA duplexes (**C**) The melting temperature (Tm) and Gibbs free energy (ΔG) of the tested compounds in DNA and RNA duplexes.

**Figure S3.**
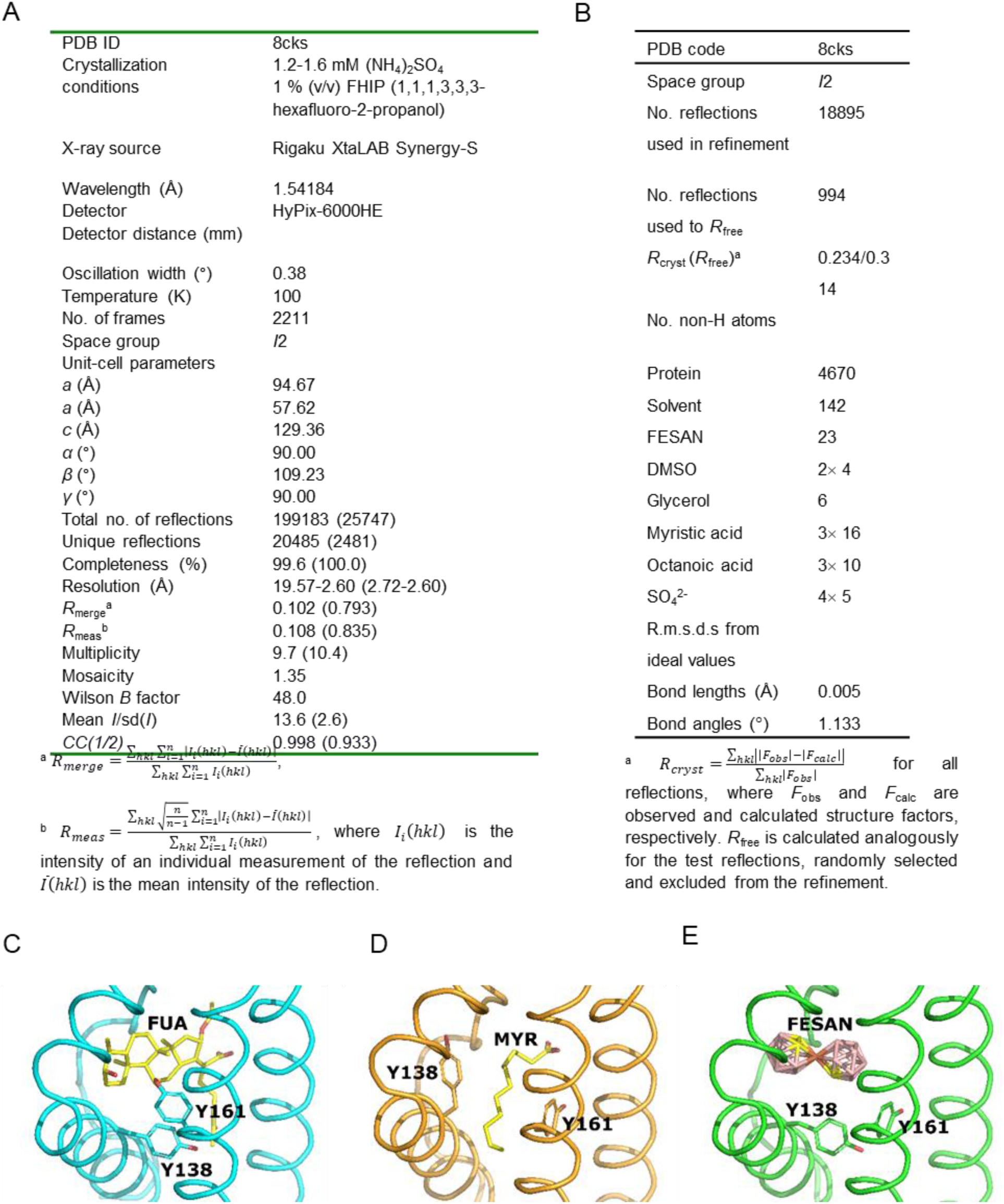
Crystal structure of human serum albumin complexed with boron cluster (FESAN). (**A**, **B**) Data collection for the HSA-FESAN crystal structure: (**C**) Comparison of the sub-domain IB structure after binding of: fusidic acid – FUA (PDB ID: 2vuf), (**D**) myristic acid - MYR (PDB ID: 1bj5;), and (**E**) FESAN (PDB ID: 8cks; present work). Ligands shown in stick representation and colored by atom type (B, salmon; C, yellow; O, red; Fe, orange).

**Figure S4.**
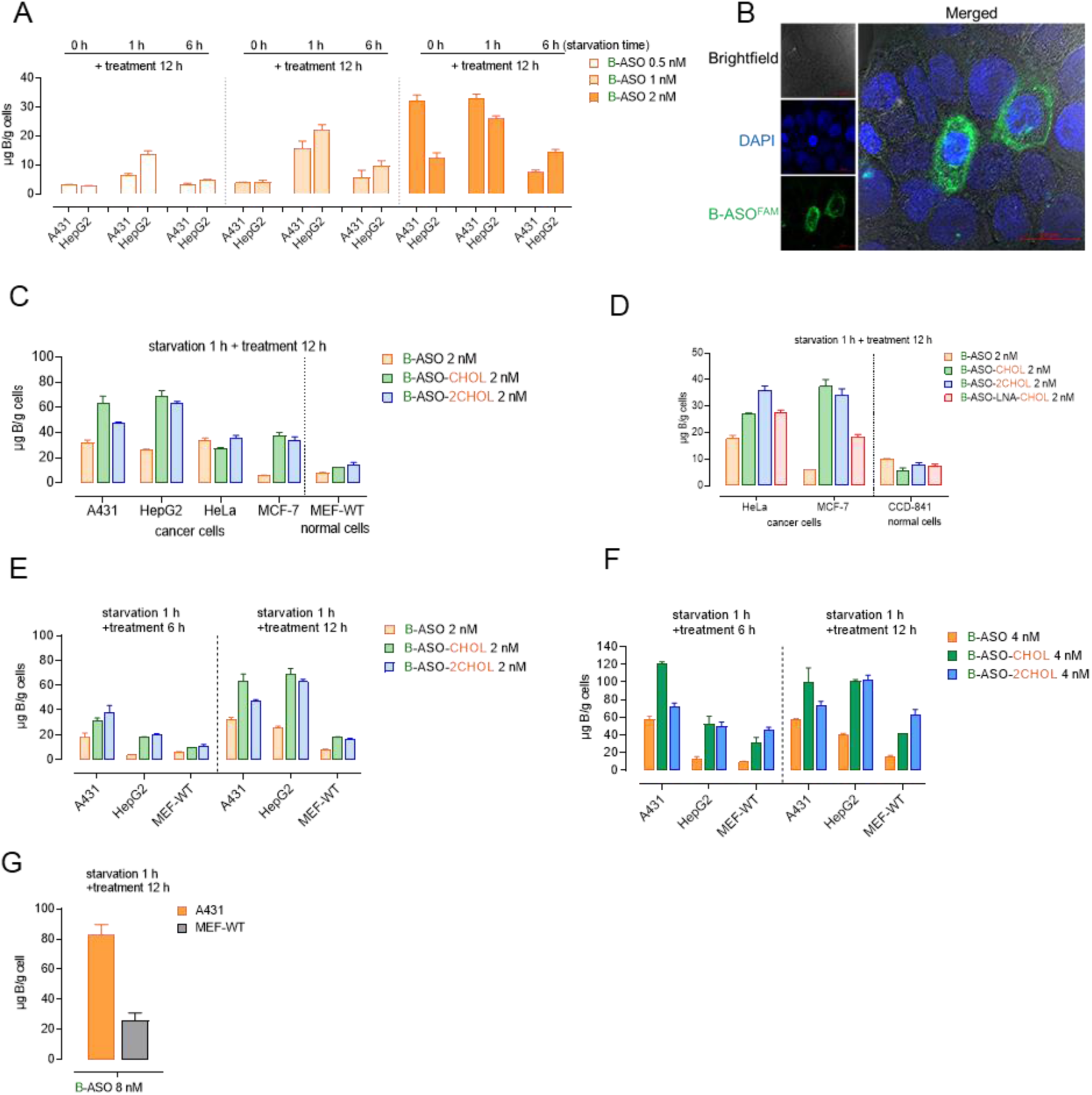
Selective uptake of B-ASO^LNA^-CHOL by cancer cells while sparing normal cells. (A) Analysis of B-ASO uptake and selectivity using various starvation durations prior to 12-h treatment; (B) Confocal microscopy images with brightfield overlay (80x) after 1-h incubation with B-ASO^FAM^ in A431 cells; (**C**, **D**) Delivery of boron atoms by B-ASO^LNA^-CHOL and its derivatives to cancer cell lines (A431, HepG2, HeLa, MCF-7) and normal cells (MEF-WT, CCD-841) measured by ICP MS. (**E**, **F**) Cellular boron saturation under different time and dose treatment (**G**) B-ASO at 8 nM selectively accumulates in A431 cancer cells than normal cells.

**Figure S5.**
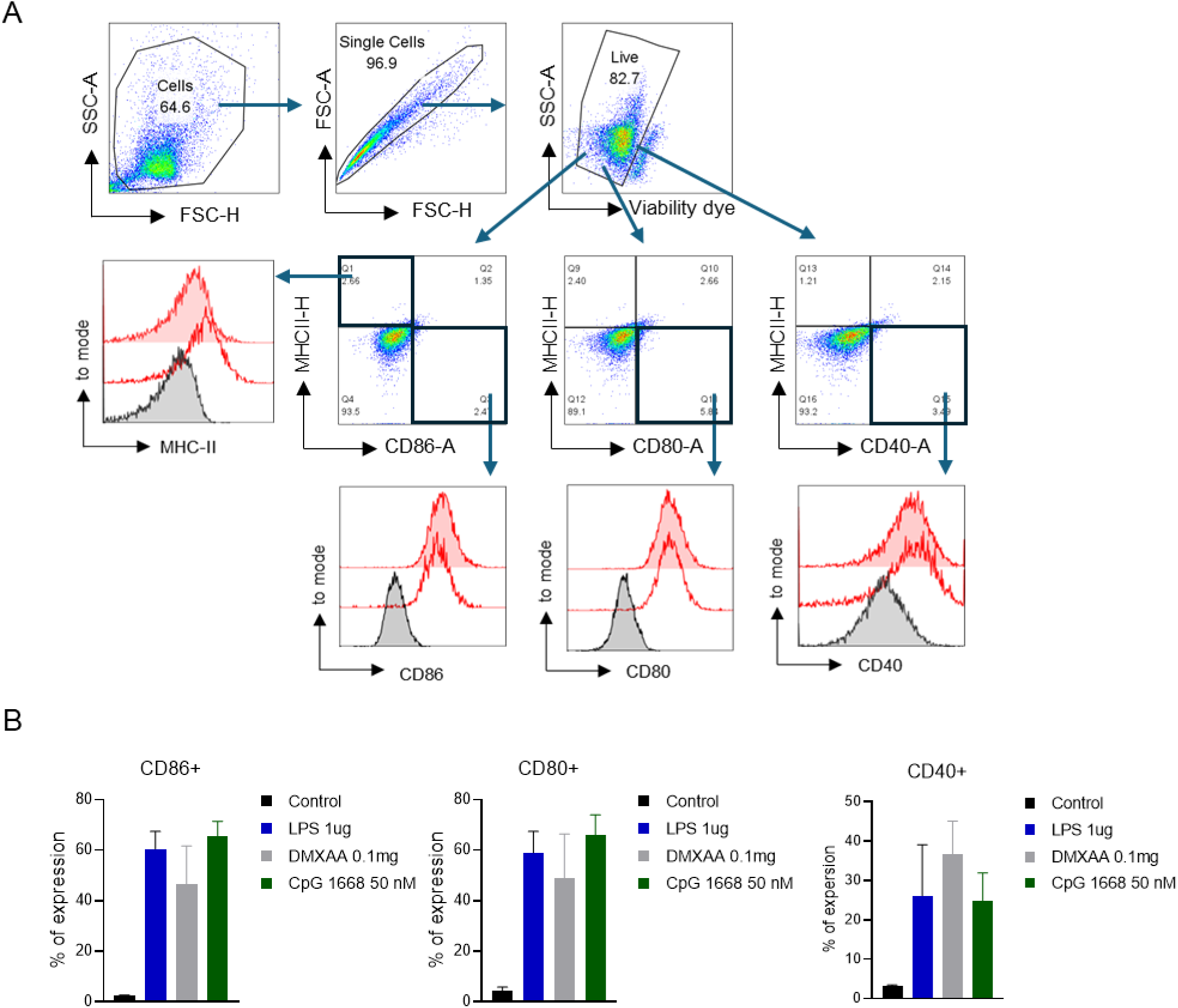
Expression of immune-related molecules on dendritic cells and macrophages. (**A**) Gating strategy for analysis of CD40, CD80, CD86, and MHC-II expression on DC2.4 mouse dendritic cells. (**B**) Expression analysis of immune markers after TLR9-mediated activation of RAW264.7 macrophages.

**Figure S6.**
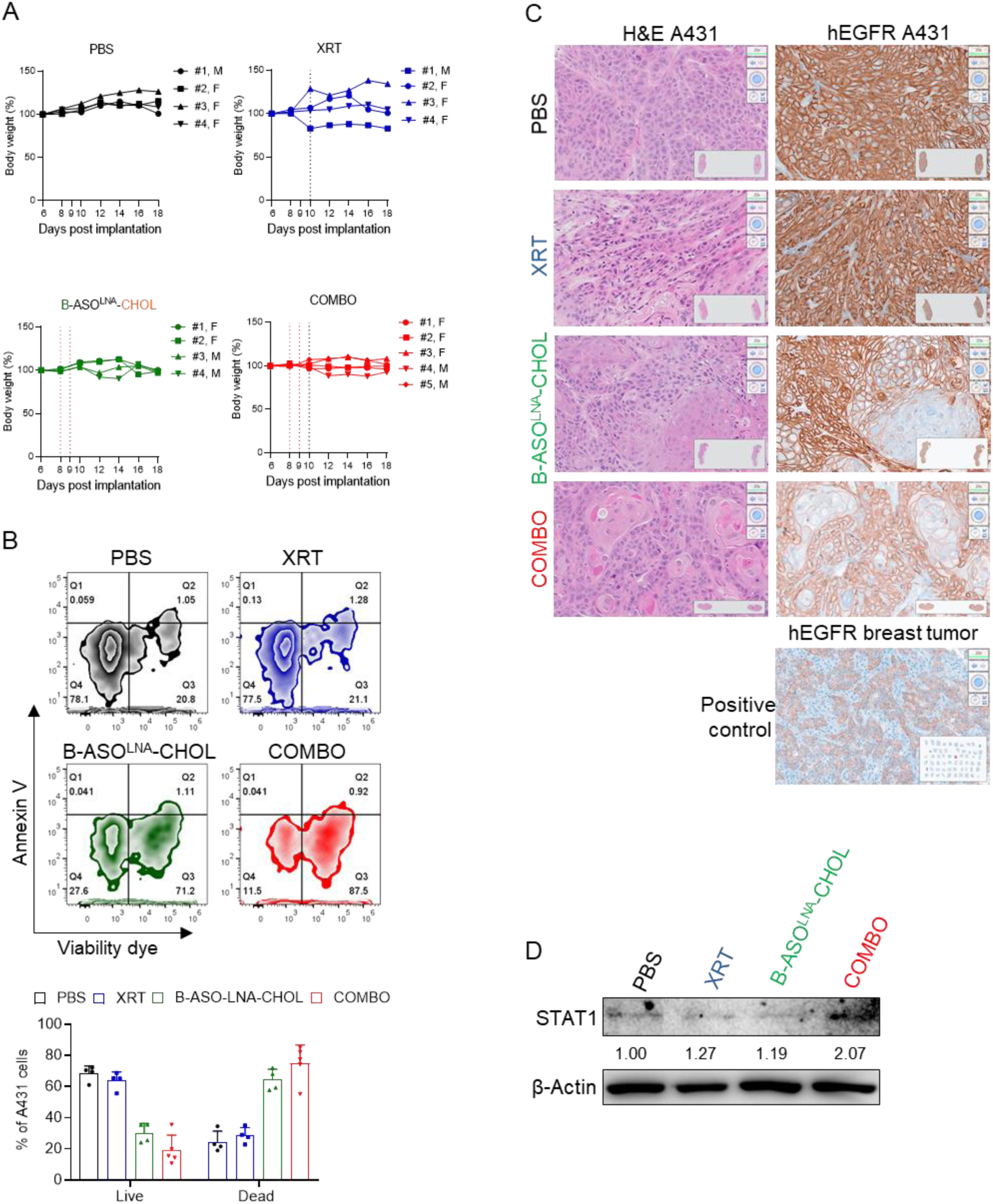
Local treatment of B-ASO^LNA^-CHOL with XRT (COMBO) significantly increased efficacy in A431 human xenografts in immunodeficient mice (NSG) compared to monotherapy. (**A**) Body weight of NSG mice monitored to 18 days after treatment; (**B**) Analysis of cell viability in the tumor using flow cytometry; (**C**) Histological H&E and anti-human EGFR immunohistochemical staining of the tumor; (**D**) STAT1 expression in the tumor analyzed by WB.

